# Monitoring phagocytic uptake of amyloid β into glial cell lysosomes in real time

**DOI:** 10.1101/2020.03.29.002857

**Authors:** Priya Prakash, Krupal P. Jethava, Nils Korte, Pablo Izquierdo, Emilia Favuzzi, Indigo Rose, Kevin Guttenplan, Sayan Dutta, Jean-Christophe Rochet, Gordon Fishell, Shane Liddelow, David Attwell, Gaurav Chopra

## Abstract

Phagocytosis by glial cells is essential to regulate brain function during development and disease. Given recent interest in using amyloid β (Aβ)-targeted antibodies as a therapy for patients with Alzheimer’s disease, removal of Aβ by phagocytosis is likely protective early in Alzheimer’s disease, but remains poorly understood. Impaired phagocytic function of glial cells surrounding Aβ plaques during later stages in Alzheimer’s disease likely contributes to worsened disease outcomes, but the underlying mechanisms of how this occurs remain unknown. We have developed a human Aβ_1-42_ analogue (Aβ^pH^) that exhibits green fluorescence upon internalization into the acidic phagosomes of cells but is non-fluorescent at physiological pH. This allowed us to image, for the first time, glial uptake of Aβ^pH^ in real time in live animals. Microglia phagocytose more Aβ^pH^ than astrocytes in culture, in brain slices and *in vivo*. Aβ^pH^ can be used to investigate the phagocytic mechanisms removing Aβ from the extracellular space, and thus could become a useful tool to study Aβ clearance at different stages of Alzheimer’s disease.

## INTRODUCTION

Glial cells make up more than half of the cells of the central nervous system (CNS) and are vital to the regulation of brain function^1^. Microglia are specialized CNS-resident macrophages that respond to pathogens and injury by clearing cell debris, misfolded protein aggregates and damaged neurons by the process of phagocytosis^2^. Mature microglia in the adult brain exhibit a ramified morphology and constantly survey their surroundings for “eat me” signals^3^ present on or released from apoptotic cells, microbes, protein deposits, dysfunctional synapses and other target substrates. After CNS injury or during neurodegenerative diseases like Alzheimer’s disease (AD) microglia become “reactive”, and change morphology, becoming rod-like or amoeboid^4^, and actively engage with their environment by secreting inflammatory cytokines like TNF-α and IL-1α. These cytokines cause functional changes in astrocytes, microglia themselves and other cells^5,6^.

During phagocytosis, proteins on the microglial cell surface, such as the Toll-Like Receptors (TLRs), Fc receptors, and scavenger receptors including CD36 and the receptor for advanced glycation end products (RAGE) among others, recognize the “eat-me” signals and engulf the target substrates into intracellular compartments called phagosomes^7–10^. The phagosomes mature by fusing with lysosomes to form highly acidic phagolysosomes and mobilize the phagocytosed material for enzymatic degradation. The pH of phagosomal organelles during this maturation process is progressively reduced^11^ from 6.0 to 4.5-4.0. Although microglia are the “professional phagocytes” of the CNS, astrocytes also competent phagocytic cells with important roles both during health, and in response to injury or in disease^12–15^. Recent evidence has demonstrated the phagocytic abilities of reactive astrocytes towards cellular debris in CNS injury^16^. Together, reactive microglia and astrocytes play a crucial role in clearing extracellular debris and cellular components and aid in remodeling the tissue environment during disease.

AD is characterized by the generation of soluble oligomers of amyloid β (Aβ) that have numerous downstream actions, including reducing cerebral blood flow^17^, inhibiting glutamate uptake that may cause hyperexcitability of neurons^18^, and inducing hyperphosphorylation of the cytoskeletal protein tau^19,20^ that leads to synaptic dysfunction and cognitive decline. Ultimately Aβ oligomers are deposited as extracellular plaques in the brain, a hallmark of AD, which contribute to neuroinflammation and neuronal death^21^. The main Aβ species generated excessively in AD is Aβ_1-42_, which is a small ~4.5 kDa peptide produced by the cleavage of amyloid precursor protein on neuronal membranes by β- and γ-secretases^22,23^. Removal of Aβ from the extracellular space by phagocytosis into microglia and astrocytes, as well as by clearance across endothelial cells into the blood or lymph vessels, is thought to limit the build-up of extracellular Aβ concentration. However, AD occurs when Aβ generation outweighs its removal^20^. Thus, to understand the onset of plaque deposition during AD (and perhaps how to prevent it) it is essential to understand glial phagocytosis and degradation of Aβ. A method that can monitor this process, especially in real time *in vivo*, will facilitate identification of the receptors that bind to Aβ and initiate its phagocytic clearance. This will allow investigation of why glia that surround Aβ plaques in AD show impaired phagocytic function^24–26^, and why in inflammatory conditions microglia may increase their phagocytic capacity depending on their state of activation^27,28^.

Current methods to study glial phagocytosis involve the use of fluorescent latex beads^29^ or particles of zymosan^30^ or *E. coli*^31^ conjugated to non-pH based fluorophores like fluorescein and rhodamine^32^. A non-pH based particle makes it hard to determine whether the particle is inside the phagosomes or outside the cell during live-cell monitoring (**Figure S1a**). Some bioparticles can be labeled with pH-sensitive dyes such as pHrodo™, however, currently available pH-sensitive dyes are not suitable for labeling disease-specific pathogenic molecules like Aβ for *in vivo* use. While non-pH sensitive fluorophore conjugates of Aβ have been used to evaluate Aβ phagocytosis^29,33^, they have several disadvantages for live-cell imaging and cannot be used for selective identification and isolation of phagocytic cells *in vivo*. First, pH-insensitive fluorophore conjugated Aβ peptides exhibit sustained fluorescence in the extracellular space (at physiological pH) thus contributing a noisy background that hinders the clear visualization of the live phagocytic cells (**Figure S1b**). Second, in live-cell imaging and in fluorescence-activated cell sorting (FACS) of live cells, it is difficult to differentiate between Aβ molecules that are internalized by the cells versus Aβ molecules that are stuck to the cell surface.

To address these issues, we have developed a pH-dependent fluorescent conjugate of human Aβ_1-42_, which we call Aβ^pH^, and characterized it using mass spectrometry, atomic force microscopy and imaging of its uptake into cells *in vitro* and *in vivo*. We show the functionality of the Aβ^pH^ probe for identifying phagocytic microglia and astrocytes in several different biological model systems such as cell lines, primary cell cultures, brain tissue slices, and *in vivo* in brain and retina. Aβ^pH^ retains an aggregation phenotype similar to that of synthetic Aβ *in vitro* and exhibits increased green fluorescence within the acidic pH range of 2.0 to 4.5 and not at the extracellular and cytoplasmic physiological pH values of 7.4 and 7.1, respectively. Aβ^pH^ can be used to visualize phagocytosis in live cells in real time without the use of any Aβ-specific antibody. It is internalized by glial cells (both astrocytes and microglia) in live rat hippocampal tissue sections *in situ*. Stereotaxic injection of Aβ^pH^ into the mouse somatosensory cortex leads to its uptake by astrocytes and microglia, following which microglia retain the Aβ^pH^ within the cells up to 3 days *in vivo* unlike astrocytes. Similarly, fast astrocyte clearance was observed in retinal tissue, using immunohistochemical staining of tissue sections. Finally, we show, for the first time, real-time phagocytosis of Aβ into microglia and astrocytes in mouse cortex *in vivo* by two-photon excitation microscopy.

## RESULTS

### Properties of a novel pH-dependent fluorescent conjugate of human Aβ_1-42_

We synthesized a new pH-sensitive fluorescent dye-labelled phagocytic Aβ probe for imaging both *in vitro* and *in vivo*, and cell sorting that allows for downstream analysis of functional subtypes of cells. We used a facile bioconjugation strategy to make our new probe safe for use with live cells *in vitro* and with live animals *in vivo*. The Aβ^pH^ conjugate was synthesized at the microgram scale by linking the synthetic human Aβ_1-42_ peptide to the amine-reactive Protonex Green 500, SE (PTXG) fluorophore (**Figure 1a**). We selected PTXG based on its ability to emit intense fluorescence at the low pH of 4.5-5.0 found within lysosomes. Conjugation of the fluorophore was performed at the side chain amine groups of the lysine residues within, and at the N-terminal of, human Aβ_1-42_ peptide. The conjugation was confirmed with matrix-assisted laser desorption/ionization-mass spectrometry (MALDI-MS) (**Figures S2, S3**) that indicates the conjugation of PTXG with the Aβ_1-42_ peptide (molecular weight >4.5K) by removal of succinimidyl ester (SE) as a leaving group. Additionally, proton nuclear magnetic resonance (^1^H-NMR) analysis of Aβ^pH^ also demonstrated the presence of PTXG as well as Aβ_1-42_ peptide (**Figures S4-S6**). The spectrum of Aβ_1-42_ peptide obtained from attenuated total reflection Fourier transform infrared spectroscopy (ATR-FTIR) also shows a strong absorption peak at 1625 cm^−1^ confirming amine functional group (**Figure S7**) and the spectrum of PTXG shows the presence of amide and ester group with absorption peaks at 1668 and 1727 cm^-1^, respectively (**Figure S8**). The conjugated product Aβ^pH^ shows a distinct peak at 1674 cm^-1^ confirming the amide bond formation between the Aβ_1-42_ peptide and PTXG dye, as expected (**Figure S9**). Collectively, these experiments confirm the formation of the peptide-dye conjugate. We also synthesized another conjugate of Aβ_1-42_ with the pHrodo-Red, NHS fluorophore (RODO) and confirmed conjugate formation from the MALDI spectrum (**Figure S10**).

**Figure 1.**
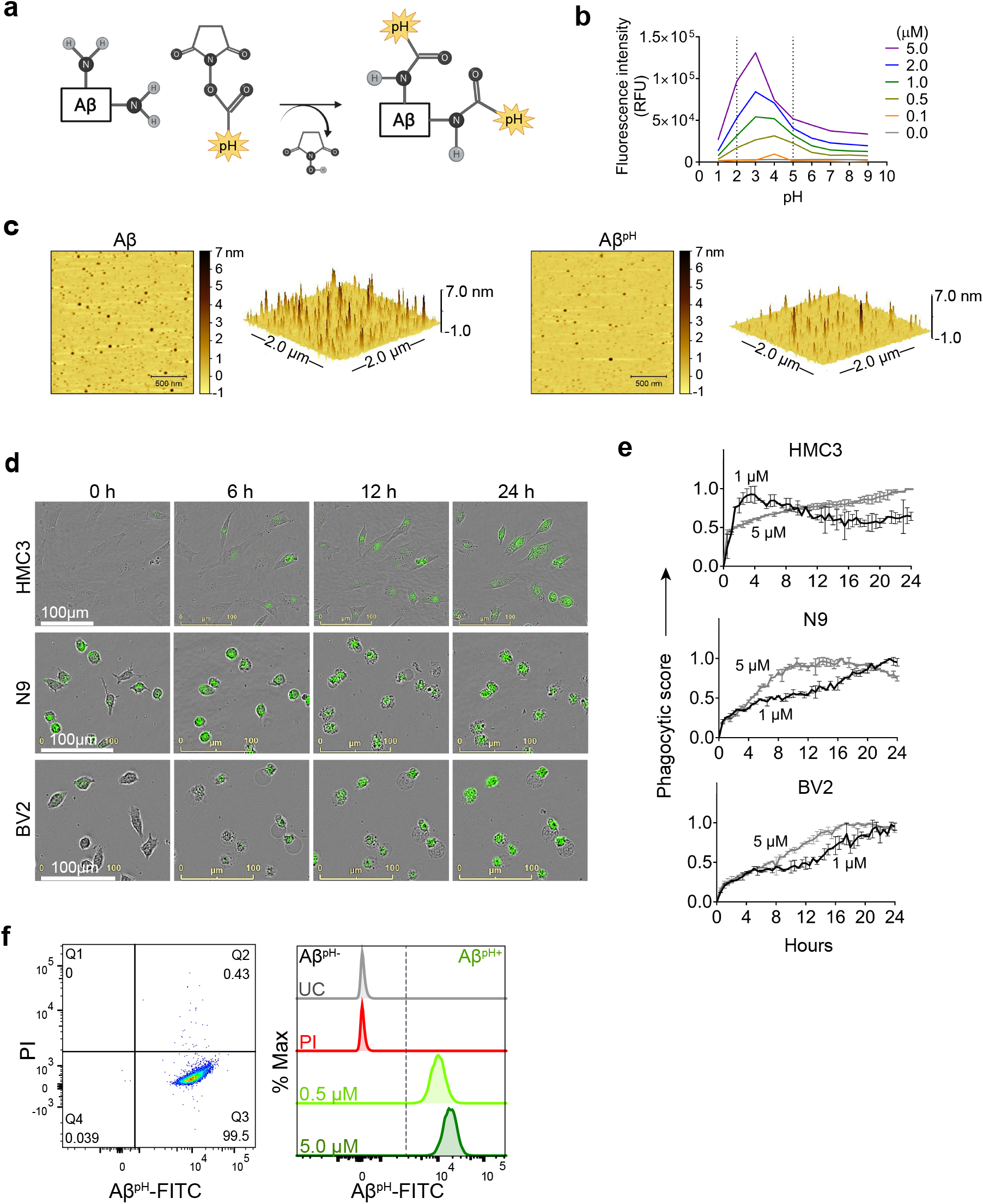
Synthesis and characterization of Aβ^pH^. **(a)** The Aβ^pH^ is synthesized by conjugating the amine-reactive pH-sensitive Protonex™ Green dye to the side chain amine groups of the lysine residues and the N-terminal of human Aβ_1-42_ peptide. **(b)** The pH-sensitivity of the Aβ^pH^ probe characterized at different concentrations from 0.1 *μ*M to 5.0 *μ*M. Increased fluorescence is observed within the narrow range of pH 2.0 to 5.0. **(c)** Atomic Force Microscopy topographic images of Aβ^pH^ oligomers compared to synthetic Aβ oligomers. Left-2D topographic image of Aβ^pH^ and synthetic Aβ oligomers. Right-3D image (2×2 *μ*m x–y). **(d)** Live cell imaging of the phagocytic uptake of 1 *μ*M Aβ^pH^ by BV2 and N9 mouse microglia and by HMC3 human microglia over 24 h. **(e)** Quantification of Aβ^pH^ phagocytic score by BV2, N9, and HMC3 microglial cells from the live cell images. **(f)** The phagocytic uptake of Aβ^pH^ by BV2 cells is measured and quantified via flow cytometry analysis. Dot plot shows live (PI^-^) and Aβ^pH+^ cells. No green fluorescence is measured in unstained cells (UC) and in dead cells stained with the PI only whereas green fluorescence is measured in cells treated with 0.5 and 5.0 *μ*M Aβ^pH^ for 1 hour (higher fluorescence is seen in cells exposed to the higher concentration of Aβ^pH^). Data shown in terms of % max, by scaling each curve to mode = 100% (y-axis).

The pH-sensitivity of the PTXG and RODO-conjugated Aβ^pH^ were confirmed by measuring their fluorescence intensities at pH 1.0 to 9.0 in solution. The PTXG-conjugated Aβ^pH^ show a sharp maximum of fluorescence between pH 5.0 and 2.0 at concentrations of 0.5, 1.0, 2.0, and 5.0 *μ*M (**Figure 1b**) with excitation/emission wavelengths of 443/505 nm. At these concentrations, the PTXG-Aβ^pH^ showed maximum fluorescence intensities between 500 and 510 nm in the 4.5-2.0 pH range (**Figure S11**). The PTXG-Aβ^pH^ conjugate exhibits low fluorescence intensity at pH values more alkaline than 5.0 including at the physiological extracellular pH of 7.4. In contrast, the RODO-conjugated Aβ^pH^ displayed increased fluorescence within a wider pH range of 1.0-7.0 for the same concentrations (**Figure S12**). Due to the broad range of fluorescence exhibited by the RODO-Aβ^pH^ conjugates, and long-term sustained fluorescence intensity of PTXG and PTXG-Aβ^pH^ (**Figure S13**), the PTXG-Aβ^pH^ conjugate was chosen for all further experiments (termed Aβ^pH^ henceforth). Lastly, we wanted to determine whether Aβ^pH^ exhibits aggregation properties similar to the aggregation of synthetic, non-conjugated Aβ. The Aβ_1-42_ and Aβ^pH^ oligomers were prepared^34^ from hexafluoroisopropanol (HFIP) treated peptide films in PBS pH 7.4 buffer at 4°C. Atomic force microscopy (AFM) revealed that the ability of Aβ^pH^ to aggregate is similar to that of the non-conjugated Aβ (**Figure 1c**) suggesting that Aβ^pH^ is suitable for biological use^35,36^.

### Aβ^pH^ uptake into human and mouse microglial cell lines

To visualize phagocytosis of Aβ^pH^ in real-time in live microglial cells, immortalized human microglial clone 3 (HMC3) cells and mouse BV2 and N9 microglial cells were treated with 1, 2 and 5.0 *μ*M concentrations of Aβ^pH^ and live-cell images were acquired every 30 minutes for 24 hours (**Figure 1d**). We observed internalization and increased fluorescence of Aβ^pH^ (implying phagocytosis) by HMC3 cells. The fluorescence was quantified as a *Phagocytic Score*, i.e. relative fluorescence compared to initial time (t=0) normalized over the 24-hour period (see **Methods**). For HMC3 cells there was an initial rapid phase of fluorescence (score) increase followed either by a slower increase in fluorescence at 5 *μ*M Aβ^pH^ concentration or a slow decrease of fluorescence from its peak value at 1 *μ*M and 2 *μ*M Aβ^pH^ concentration (**Figures 1e, S14**). This suggests rapid initial uptake of Aβ^pH^, followed by intracellular degradation of Aβ^pH^ which occurs either more rapidly than the influx (giving a slow decline) or less rapidly than the influx (giving a slowed increase) (**Figure S14**). Cells that did not phagocytose Aβ^pH^ did not display any green fluorescence thereby differentiating Aβ^pH^-specific phagocytic and non-phagocytic microglial cells in real time. Rodent microglial cell lines (BV2 and N9) showed a peak of *Phagocytic Score* at 12-16 hours for N9 and 16-20 hours for BV2 at 5 *μ*M Aβ^pH^ treatment, compared to the HMC3 human microglial cell line that showed a gradual increase in phagocytosis over the 24-hour treatment period for the same concentration. Interestingly, for the lower Aβ^pH^ doses of 1 *μ*M and 2 *μ*M, the peak value of *Phagocytic Score* for HMC3 cells was within the initial 4 hours compared to the gradual increase for the rodent cell lines over the 24 hour period (**Figure S14**). Using live-cell imaging, we also observed interesting morphological differences over time between phagocytic and non-phagocytic microglial cells. During the initial 2 hours, many cells displayed an elongated, branched morphology followed by an amoeboid morphology during subsequent time points with increased fluorescence (**Movies S1-3**). Thus, the Aβ^pH^ reporter can be used to visualize Aβ-specific phagocytosis in real-time and can be used in experiments to evaluate the enhancement or inhibition of microglial phagocytosis for *in vitro* screening of drug candidates for AD.

### Flow cytometry of Aβ^pH^ phagocytic cells and staining of fixed Aβ^pH^ after phagocytosis by primary cultured microglia and astrocytes

We next determined whether Aβ^pH^ can be used to purify phagocytosing cells using fluorescent activated cell sorting (FACS). By adding Aβ^pH^ to BV2 cells *in vitro*, we show that live microglial cells that phagocytose Aβ^pH^ can be easily analyzed with flow cytometry without the need for traditional dyes or antibodies to detect Aβ (**Figure 1f**). Phagocytic uptake of Aβ^pH^ by BV2 microglia was evident with a green fluorescence peak within live cells when the cells were treated with 0.5 *μ*M and 5.0 *μ*M Aβ^pH^ for 1 hour in culture (**Figures 1f, S15**), with unstained and live/dead stained cells as controls. There was a slight increase in Aβ^pH^ fluorescence at 1 hour for a 10-fold increase in Aβ^pH^ concentration for BV2 cells. The green fluorescence signal indicates internalization of Aβ^pH^ into the cellular acidic organelles thereby avoiding detection of peptide sticking to the cell surface. Thus, Aβ^pH^ fluorescence after phagocytosis is sufficiently bright to enable FACS experiments, enabling single cell analysis of Aβ^pH+^ phagocytic and Aβ^pH-^ non-phagocytic microglia and other glial cells.

Fixing cells that have internalized Aβ^pH^, and using specific antibodies and dyes to study specific molecular processes, can help identify molecular mechanisms involved during Aβ phagocytosis. To determine whether Aβ^pH^ can maintain its fluorescence in fixed cells, we used primary mouse CD11b^+^ microglial cells isolated from 3-5 month old mice and cultured in reduced-serum TIC medium^37^ as well as BV2, N9, and HMC3 microglia. After Aβ^pH^ (5.0 *μ*M) treatment for 2 hours, the cells were fixed in 4% paraformaldehyde followed by addition of red phalloidin dye (a podosome core marker for F-actin) to visualize actin filaments and the cell body with confocal microscopy (**Figure S17, S18**). Confocal imaging of the fixed Aβ^pH^-treated microglia showed green fluorescence within the red actin filaments thereby confirming the uptake of Aβ^pH^ peptides by the cells. Further, LysoTracker DND-99 was used to confirm localization of Aβ^pH^ within acidic organelles such as phagosomes or phagolysosomes. The co-localization of green fluorescent Aβ^pH^ along with the red signal from the LysoTracker dye confirmed the presence of Aβ^pH^ within acidic phagolysosomes after 2 hours in HMC3, N9, and BV2 microglia cell lines (**Figures 2a, S19**). Similarly, uptake of Aβ^pH^ into intracellular acidic organelles was also confirmed in CD11b^+^ primary microglia that were isolated from adult mice and cultured in low serum containing defined media (**Figure 2b**). Finally, we also detected the green fluorescence signal within CD11b^+^ primary microglia (Aβ^pH+^ cells) at 0.5, 1.0, and 2.0 *μ*M Aβ^pH^ concentrations via flow cytometry (**Figures 2c, S16**) after 1 hour of treatment, and found an increase in fluorescence with Aβ^pH^ concentration. The unstained and live/dead stained cells were used as controls. Microglia recognize and phagocytose Aβ peptides through scavenger receptors such as^38–40^ Toll-Like Receptor 2 (TLR2) Cluster of Differentiation 14 (CD14) and Triggering Receptors Expressed on Myeloid Cells 2 (TREM2). Thus, the Aβ^pH^ reporter that we have developed could serve as a valuable chemical tool to delineate the role of receptor proteins involved in Aβ uptake by glial cells, using specific antibodies or CRISPR-Cas9 genetic screens.

**Figure 2.**
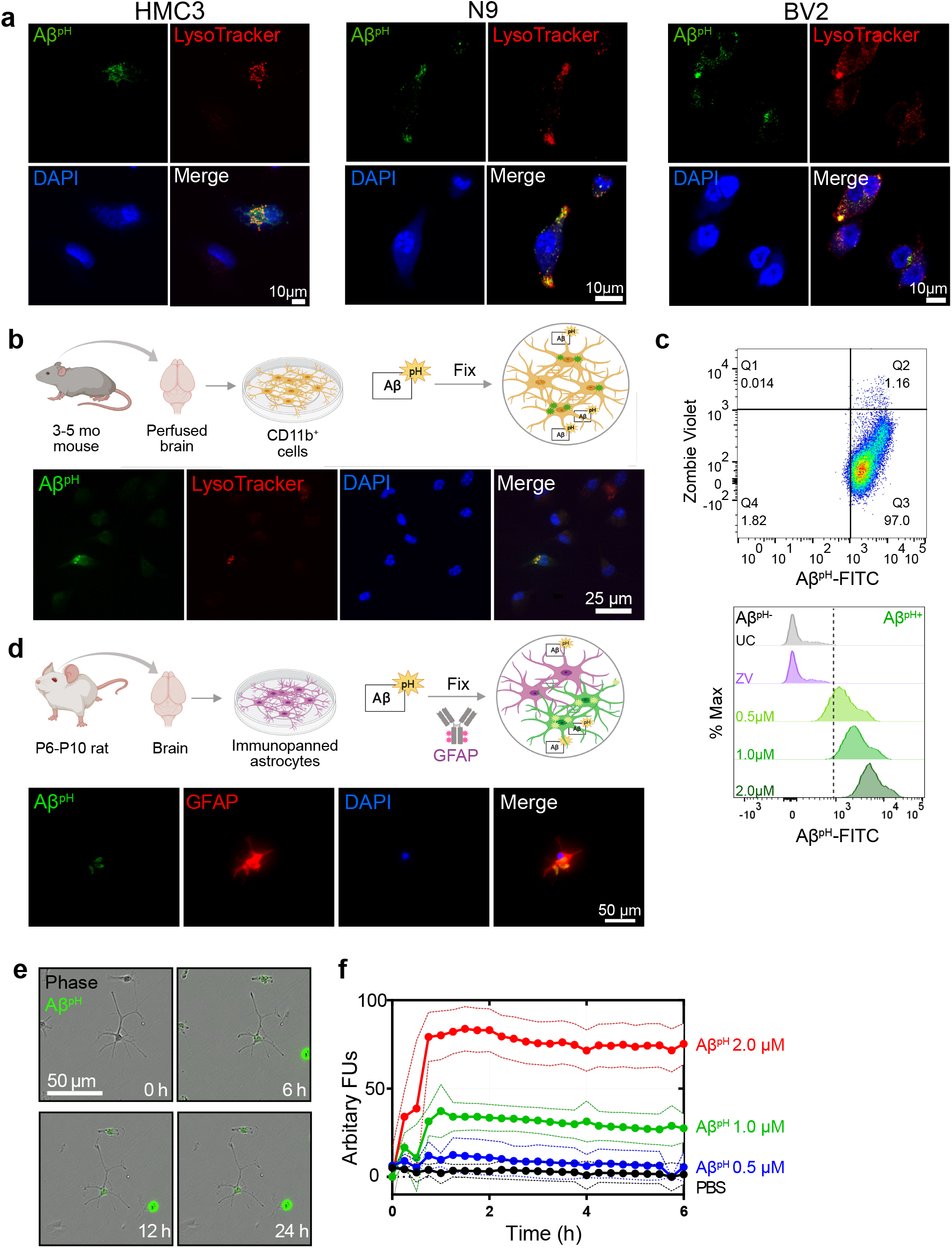
Fluorescence of internalized Aβ^pH^ is retained in fixed cells. **(a)** Confocal images of fixed HMC3, N9, and BV2 cells showing the uptake of Aβ^pH^ (green). Cells are stained for acidic intracellular organelles (LysoTracker Red, confirming co-localization of the Aβ^pH^ within the acidic intracellular organelles) and nuclei (DAPI, blue). No antibody is required to detect Aβ^pH^. **(b)** Primary mouse microglia grown in defined, reduced-serum media phagocytose Aβ^pH^ *ex vivo*. Cells are fixed and stained for nuclei and show Aβ^pH^ colocalized in the acidic organelles with LysoTracker Red. **(c)** The phagocytic uptake of Aβ^pH^ by primary microglia is measured and quantified via flow cytometry analysis. Dot plot shows live (ZV^-^) and Aβ^pH+^ cells. No green fluorescence is measured in unstained cells (UC) or dead cells stained with the ZV live/dead stain only whereas green fluorescence is measured in cells treated with 0.5, 1.0, and 2.0 *μ*M Aβ^pH^ for 1 hour. Data shown in terms of % max, by scaling each curve to mode = 100% (y-axis). **(d)** Primary immunopanned rat astrocytes also phagocytose Aβ^pH^ in serum-free conditions. Cells are fixed and stained for astrocyte specific GFAP antibody (red) and nuclei. **(e)** Uptake of Aβ^pH^ over time by primary immunopanned astrocytes as observed in live cells in real time. **(f)** Quantification of uptake of 0.5, 1.0, and 2.0 *μ*M Aβ^pH^ by primary astrocytes.

In addition to microglia, astrocytes have been shown to exhibit phagocytic characteristics after ischemic injury^16^ and can phagocytose^29^ extracellular Aβ. Therefore, we tested whether primary immunopanned cultured mouse astrocytes phagocytosed Aβ^pH^. Indeed, this was the case, with retention of the phagocytic Aβ^pH^ signal after methanol-fixation and staining of cells with GFAP antibody (**Figure 2d**). Aβ^pH^ phagocytosis could also be detected with live cell imaging (**Figure 2e**). Quantification of Aβ^pH^ uptake at different concentrations showed internalization increasing for approximately 1 hour, followed by a sustained fluorescence within the cells which may reflect a balance between phagocytosis and degradation of the probe (**Figure 2f**). Thus, Aβ^pH^ may be a viable candidate for concentration and time-dependent studies of glial cell phagocytosis *in vitro*. The sustained fluorescence of Aβ^pH^ seen for up to 6 hours inside cultured microglia and astrocytes suggests it is chemically stable under physiological conditions – allowing for long-term use *in vivo*.

### Aβ^pH^ uptake by microglia and astrocytes in hippocampus, cortex, and retina

To assess phagocytic uptake of Aβ^pH^ in the hippocampus, a brain area that is crucial for learning and memory, we applied 5 *μ*M Aβ^pH^ for 1.5 hours at 37°C to live hippocampal slices from postnatal day (P)12 rat. The tissue slices were then fixed and stained with glial cell specific antibodies (**Figure 3a**). Microglia phagocytosed Aβ^pH^ as seen by the localization of Aβ^pH^ within the IBA1^+^ myeloid cells in these tissues (**Figure 3b**). Our experiments revealed green fluorescent signal both within IBA1^+^ microglial cells as well as outside the IBA1^+^ cells (**Figure S20**), presumably reflecting phagocytic uptake of Aβ^pH^ by cells other than microglia, such as astrocytes. Indeed, staining with GFAP antibody demonstrated internalization of Aβ^pH^ within GFAP^+^ astrocytes (**Figure 3c**). Summing over cells, approximately 60% of the taken up Aβ^pH^ was phagocytosed by microglia and 40% by astrocytes (**Figure 3d**). Interestingly, in astrocytes the phagocytosed Aβ^pH^ was distributed more homogeneously throughout the GFAP area than the phagocytosed Aβ^pH^ in IBA1 stained microglia. This is expected since in microglia the acidic compartments (where Aβ^pH^ is present) are mostly juxtanuclear whereas IBA1 is known to be enriched in podosomes and podonuts^41^. In contrast, in astrocytes the acidic compartments occur all over the cell body^42^.

**Figure 3.**
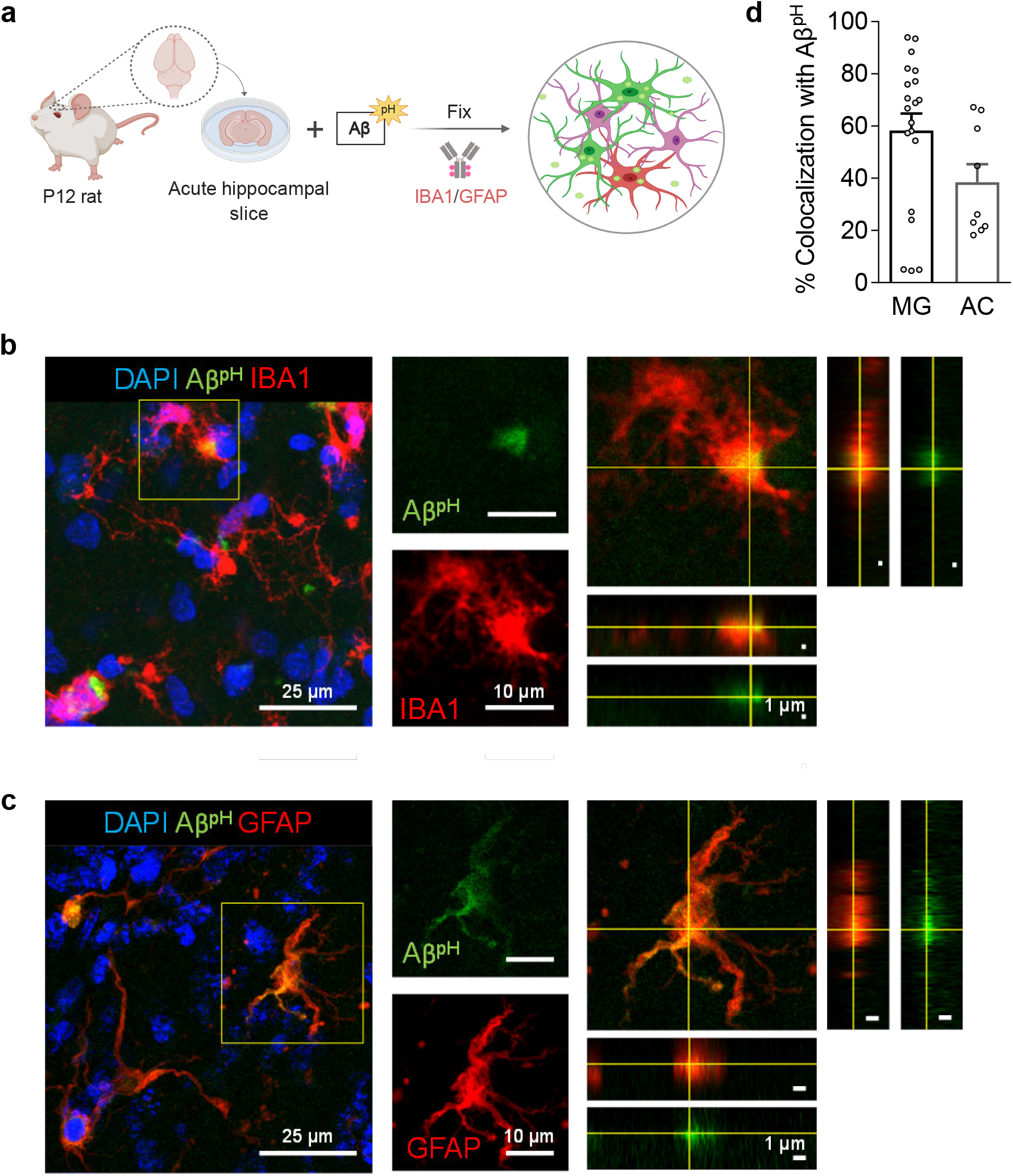
Aβ^pH^ is phagocytosed by both microglia and astrocytes *in situ* in rat hippocampal tissue sections. **(a)** Schematic of phagocytosis assay in rat hippocampal tissue slices. **(b)** Representative 2D maximum projection of a confocal z-stack showing microglia phagocytosing Aβ^pH^. Closeup of the indicated cell (yellow square) is shown on the right. Orthogonal projections at the level of the crosshairs show internalization of Aβ^pH^ within the microglia. **(c)** Representative 2D maximum projection of a confocal z-stack showing astrocytes phagocytosing Aβ^pH^. Closeup of the indicated cell (yellow square) is shown on the right. Orthogonal projections at the level of the crosshairs show internalization of Aβ^pH^ within the astrocyte. **(d)** Quantification of Aβ^pH^ colocalized with microglia and astrocytes, as defined by IBA1^+^ and GFAP^+^ staining, respectively. Data shown as mean±s.e.m. collected from 20 and 8 slices for microglia and astrocytes respectively (from 3 animals).

We next tested the Aβ^pH^ probe in different *in vivo* settings. After intracranial injection of Aβ^pH^ into the somatosensory cortex of wild-type P7 C57BL/6J mice (**Figure 4a**), phagocytosis of the Aβ^pH^ by IBA1^+^ microglia and GFAP^+^ astrocytes was assessed by fixing the corresponding tissue sections at 24 and 72 h after injection. At 24 h, Aβ^pH^ was observed in the injected area and appeared to be enclosed within cell bodies. There was less astrocyte uptake of Aβ^pH^ compared to microglia, which may be a result of a reactive response by astrocytes which has been shown to downregulate phagocytic pathways in some states^5^ (**Figure 4b, d**). At 72 hours after the Aβ^pH^ injection there was also Aβ^pH^ visible in the pial surface and in the periventricular white matter. With increased magnification, we observed cell bodies containing Aβ^pH^ that were positive for the microglial (IBA1) marker (**Figure 4c**) but astrocytes (GFAP^+^) localized at the injection site showed very little Aβ^pH^ signal (**Figure 4d**). Thus, microglia engulf the majority of the Aβ^pH^ under these conditions *in vivo* with astrocytes contributing also at early times.

**Figure 4.**
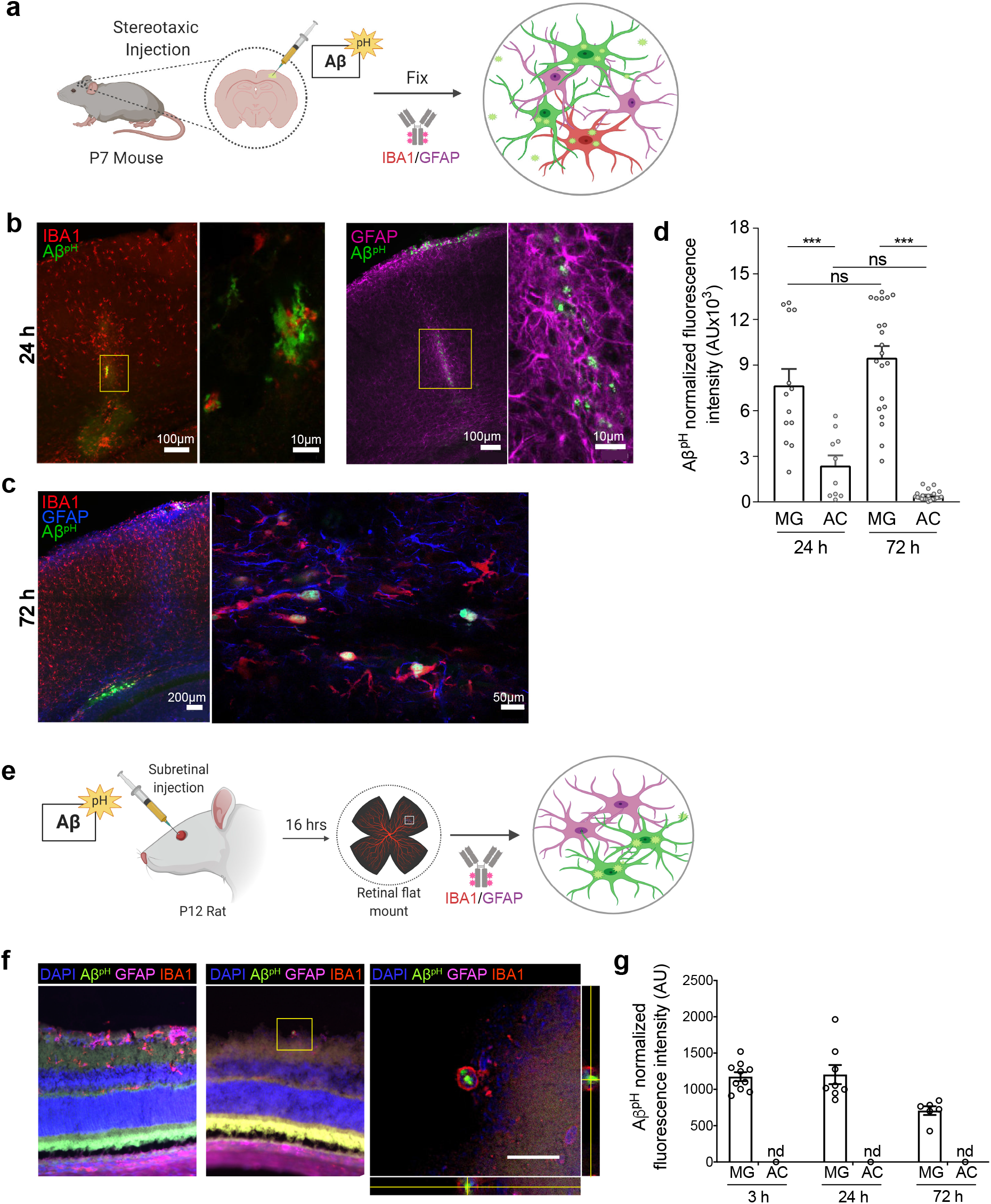
Aβ^pH^ is phagocytosed by cortical microglia and astrocytes *in vivo* and by rat retinal astrocytes *in vivo*. **(a)** Schematic of stereotaxic microinjection of Aβ^pH^ in the somatosensory cortex of P7 mouse followed by staining of fixed tissue section after 24 and 72 hours. **(b)** Phagocytic uptake of Aβ^pH^ by IBA1^+^ microglia and GFAP^+^ astrocytes in the periventricular white matter at the 24 h timepoint. **(c)** IBA1^+^ microglia show bright green fluorescence at 72 h in the same region indicating presence of Aβ^pH^ within the cells at this timepoint. GFAP^+^ astrocytes do not show any green fluorescence in this region at this timepoint suggesting either degradation of the peptide or insufficient Aβ^pH^ concentration for detectable phagocytic uptake by these cells. **(d)** Quantification of Aβ^pH^ fluorescence within microglia and astrocytes located in the pia and white matter regions show more Aβ^pH^ uptake by microglia compared to astrocytes. **(e)** Schematic of subretinal injection of Aβ^pH^ to evaluate its *in vivo* uptake by retinal microglia and astrocytes. **(f)** GFAP^+^ rat retinal astrocytes phagocytose Aβ^pH^ *in vivo*. **(g)** Quantification of Aβ^pH^ uptake into retinal IBA1^+^ microglia and GFAP^+^ astrocytes. No fluorescence was detected in astrocytes at these 3 time points (n.d.). Data shown as mean±s.e.m. from 2 animals.

Next, we injected Aβ^pH^ into the vitreous of the eye in postnatal rats to eliminate possible complications due to glial scarring resulting from cortical injection, and low penetration through the blood-brain barrier as a result of peripheral delivery (**Figure 4e**). We injected 1 *μ*l of Aβ^pH^ intravitreally and left the animals for 3, 24, or 72 hours. At the end of the experiment, retinae were removed, fixed in 4% paraformaldehyde, and whole mount retinal preparations were made. Astrocytes and microglia in the retinal ganglion cell layer were labelled with antibodies to GFAP and IBA1 respectively, and co-localization with the fluorescent signal from the injected Aβ^pH^ probe was determined. We observed that IBA1+ microglia contained Aβ^pH^ at all time points, but did not detect any Aβ^pH^ signal in GFAP^+^ astrocytes (**Figure 4f-g**). The Aβ^pH^ positive microglia were less in the retina compared to other brain regions, suggesting that clearance from the eye was more rapid than clearance from the brain parenchyma in other experiments (above).

The *in vivo* investigation of Aβ phagocytosis described above consists of imaging fixed tissue sections with confocal microscopy. To date, phagocytosis of Aβ has not been observed in live animals in real time. Using Aβ^pH^, we observed phagocytic uptake in live rodents in real time using two-photon microscopy (**Movies S5, S6**). Here, 5.0 *μ*M Aβ^pH^ was applied onto the cortical surface of live mice through a cranial window for 10 minutes during two-photon imaging in the barrel cortex (**Figure 5a**). This showed an increase in green fluorescence in the cell somata after Aβ^pH^ application, indicating phagocytic uptake of Aβ^pH^ (**Figure 5b**). Quantification of the fluorescence within the cell somata showed rapid Aβ^pH^ uptake after the peptide was added to the cranial window followed by stabilization of the signal at around 20 minutes (**Figure 5c**). The brains were later fixed by cardiac paraformaldehyde perfusion and stained with antibodies to identify the cell types mediating the Aβ^pH^ uptake. Phagocytosed green Aβ^pH^ was present in microglia labeled for IBA1 and colocalized with the microglia/macrophage lysosomal protein CD68, and Aβ^pH^ was also seen in GFAP^+^ astrocytes (**Figure 5d**). Integrating the fluorescence over the different classes of labeled cells showed that about 70% of the phagocytosed Aβ^pH^ was taken up by microglia and about 25% by astrocytes (**Figure 5e**). These percentages are approximately similar to those seen in the brain slice experiments of Fig. 3d. Thus, microglial uptake of Aβ^pH^ dominates over astrocyte uptake.

**Figure 5.**
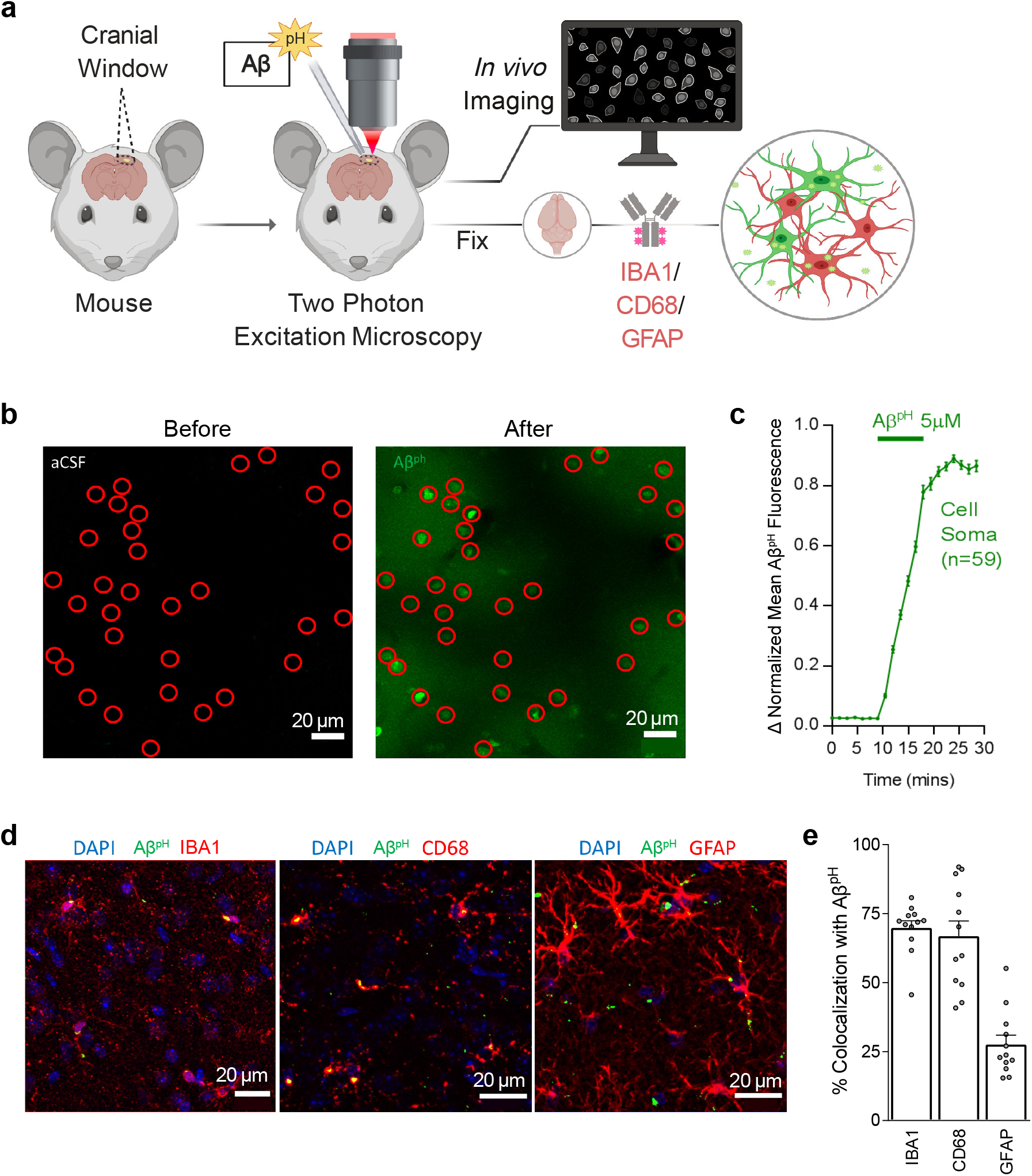
Aβ^pH^ is phagocytosed by microglia and astrocytes *in vivo* in the cerebral cortex. **(a)** Schematic of how Aβ^pH^ phagocytic uptake is imaged through a cranial window *in vivo* in real time using two-photon excitation microscopy. **(b)** *In vivo* two-photon imaging of the mouse barrel cortex before and after topical application of Aβ^pH^. The fluorescence increases in cell somata (indicated by red circles) reflecting Aβ^pH^ uptake. **(c)** Quantification of mean Aβ^pH^ fluorescence in cell somata over time. The data were normalized to the maximum mean Aβ^pH^ fluorescence for each cell and then averaged. Data shown as mean±s.e.m. N=59 somata from 2 animals. **(d)** 1.5 to 3 hours after *in vivo* two-photon imaging of Aβ^pH^, animals were perfusion-fixed and cortical slices were stained for microglia, microglia lysosomes/endosomes and astrocytes using IBA1, CD68, and GFAP antibodies, respectively. **(e)** Quantification of Aβ^pH^ colocalization with IBA1, CD68, and GFAP suggests that most Aβ^pH^ is taken up by microglia and astrocytes *in vivo*. Data shown as mean±s.e.m. N=12 stacks from 3 animals.

## DISCUSSION

The phagocytic capacity of glial cells has been measured using traditional dyes and fluorophore-labeled particles such as fluorescein or rhodamine-coated *E. coli* and beads, however, these experiments entail some major drawbacks: (1) difficulty in differentiating between adherent versus internalized particles, (2) use of additional reagents (like trypan blue or ethidium bromide) and additional experimental steps required to quench the extracellular fluorescence prior to analysis by flow cytometry and imaging^43^, (3) quenching of the fluorescence within the acidic environment of the phagosomes, and (4) lack of specificity of the neuronal target substrate (beads, *E. coli*, zymosan, etc.)^44^. Here, we have synthesized a pH-sensitive fluorogenic Aβ reporter, Aβ^pH^, at a microgram scale and characterized the reporter in detail (**Figures 1a-c, S2-13**). We demonstrated the functionality of Aβ^pH^ in mouse and human microglial cell lines, in *in vitro* cultures of primary microglia and astrocytes, in acute hippocampal slices from mouse brains, in mice *in vivo* using stereotaxic injections followed by fixation, and in live mice *in vivo*. The Aβ^pH^ reporter is a useful tool for answering questions related to mechanisms of Aβ phagocytosis and cellular inflammatory responses, and for developing therapeutic strategies to promote Aβ clearance.

We observed phagocytic uptake of Aβ^pH^ in three different microglial cell lines and primary glial cell cultures in real-time (**Figures 1d-e, 2e-f**) and using confocal imaging (**Figure 2a-b, d**). We show the utility of Aβ^pH^ to be used without the need for any antibodies to identify or detect Aβ using flow cytometry (**Figures 1f, 2c**) which will greatly benefit experimental design and outcome by reducing the number of steps required in assays. While microglia cultured *in vitro* and *ex vivo* do not completely recapitulate the transcriptomes of microglia *in vivo*^45^, microglial cell culture models provide a convenient system for screening chemical and biological molecules in a rapid and high-throughput manner. We also demonstrate the utility of Aβ^pH^ in functional assays to evaluate cellular phagocytosis. Further, we demonstrated the phagocytic uptake of Aβ^pH^ in live tissue hippocampal slices that preserve the 3-dimensional nature of the cell microenvironment and can serve as an additional useful model to study Aβ-related biology in a tissue environment. Microglia were more efficient in internalizing Aβ^pH^ than astrocytes (**Figures 3a-d**). During the period of this assay it is unlikely that astrocytes would have become “reactive” so it appears that Aβ clearance is not just a property of reactive astrocytes^46–48^, and astrocytes that are most closely associated with neurodegeneration appear to be of a reactive phenotype that downregulates phagocytic pathways^5^, suggesting that the engulfment of Aβ^pH^ by astrocytes is a normal physiological function of these cells, though this may change with disease progression.

Most importantly, we demonstrated the utility of using of Aβ^pH^ in various *in vivo* models. Stereotaxic injection of Aβ^pH^ into the mouse somatosensory cortex was followed by uptake by microglia and astrocytes (**Figure 4a-d**), injecting Aβ^pH^ into the vitreous of the eye was followed by uptake by retinal microglia (**Figure 4e-g**), and the real-time uptake of Aβ by microglia and astrocytes was demonstrated *in vivo* in live mice using two-photon microscopy (**Figures 5a-c**). We confirmed the identity of the cells observed with two-photon excitation by using IBA1 to label microglia, CD68 (a phagocytosis-specific marker) to label microglia and macrophages, and GFAP to label astrocytes (**Figures 5d-e**). The time course of uptake *in vivo* was rapid, with phagocytosed Aβ^pH^ fluorescence reaching a peak level within 20 mins (this may reflect a balance between continued phagocytosis and intracellular degradation). Approximately two thirds of the Aβ^pH^ was phagocytosed by microglia and one third by astrocytes, again on a time scale too rapid for astrocytes to have become reactive.

The development of Aβ^pH^, in particular the ease with which it can be produced in the lab, will facilitate the characterization of different populations of cells that remove (or do not remove) Aβ by phagocytosis in various conditions, including AD, trauma-induced amyloidopathy and Down’s syndrome. Combining this tracer with transgenic labeling of microglia or other cell types (e.g. *Tmem119*-tdTomato mice^49^ that exhibit red fluorescence in microglia) will allow the transcriptome and proteome of different subpopulations of heterogeneous phagocytic cells to be defined (including young and old cells^50–52^, and male and female cells^53^), facilitating the discovery of new mechanisms, targets and functional biomarkers. Understanding the clearance of Aβ is fundamental to understanding the onset of AD^54,55^, and having a quantitative technique to assess Aβ phagocytosis should contribute significantly to this. We expect that Aβ^pH^ will become a useful tool to facilitate the study of similarities and differences between phagocytic and non-phagocytic cell mechanisms *in vivo* during health and disease.

## METHODS

### Animal Ethics

Animal maintenance and isolation of primary microglia were performed according to Purdue Animal Care and Use Committee guidelines and approval (protocol number 1812001834). Injection of Aβ^pH^ into the retina of rats were completed in accordance with the National Institute of Health Stanford University’s Administrative Panel on Laboratory Animal Care. Purification of rat primary astrocyte was completed in accordance with NYU Langone School of Medicine’s Institutional Animal Care and Use Committee (IACUC) guidelines and approval. All rats were housed with *ad libitum* food and water in a 12 hour light/dark cycle. Standard Sprague Dawley rats (Charles River, #400) were used in all retinal experiments. Animal maintenance and experimental procedure for the intracranial stereotaxic injections of Aβ^pH^ in mice were performed according to the guidelines established by the IACUC at Harvard Medical School (protocol number IS00001269). Experiments on brain slices and *in vivo* 2-photon imaged mice were carried out under a UK government licence (PPL 70/8976, awarded after local ethical review and UK Home Office assessment) to David Attwell, in accordance with all relevant animal legislation in the UK.

### Synthesis of pH-sensitive fluorescent human Aβ conjugate with Protonex Green™ (Aβ^pH^)

Human amyloid-beta (Aβ_1-42_) was purchased from AnaSpec., Inc (Cat. #AS-20276), Protonex™ Green 500 SE was from AAT Bioquest, Inc. (Cat. #21215), and pHrodo-Red SE was from Thermo Fisher Scientific (Cat. #P36600). To 200 *μ*l aliquot of Aβ_1-42_ (1 mg/ml in 1M NaHCO_3_, pH ~8.5) was added 10 equivalents of Protonex-Green 500 (PTXG), SE dye (88 *μ*l from 5 mM anhydrous stock in DMSO) and incubated at room temperature for 3 hours in the dark (vial wrapped in aluminum foil: note: add 100 *μ*l ultrapure water if the solution becomes viscous). The additional 5 equivalents of PTXG, SE dye (44 *μ*l from 5 mM DMSO stock) was added and incubated under the same conditions for 3 hours. The crude reaction mixture was diluted with 1 ml ultrapure water and the conjugated product was dialyzed by Pierce Protein Concentrators [PES, 3K Molecular Weight Cut-Off (MWCO); Thermo Fisher Scientific, Cat # PI88514] at 4500 *g* for 30-45 minutes in a swinging bucket centrifuge to remove the small molecular weight fragments. The concentrated solution was diluted with 0.5 ml ultrapure water and dialyzed again for 30 minutes as done previously. Finally, MALDI-MS spectrum was recorded to confirm the conjugation of PTXG with Aβ_1-42_.

### Synthesis of pH-sensitive fluorescent human Aβ_1-42_ conjugate with pHrodo (RODO-Aβ^pH^)

A solution containing Aβ_1-42_ (570 *μ*g, 12.63 nmol) was prepared in 1M NaHCO_3_ (pH 8.5, 570 *μ*l) and the pHrodo Red-NHS (1 mg) stock solution was prepared in anhydrous DMSO (150 *μ*l) (~10.2 mM). Next, the Aβ solution (570 *μ*l) and pHrodo stock solution (0.6314 *μ*mol, 61.5 *μ*l of stock solution) were mixed and incubated at room temperature for 6 hours while wrapped with aluminum foil. This crude reaction mixture was then diluted with ultrapure water (1 ml) and the conjugated product was dialyzed by Pierce Protein Concentrators (PES, 3K MWCO) at 4500 g for 30-45 minutes in a swinging bucket centrifuge to remove the small molecular weight fragments. The concentrated solution was diluted with ultrapure water (0.5 ml) and dialyzed again for 30 minutes as before. Finally, the MALDI-MS spectrum was recorded to confirm the conjugation of pHrodo-Red fluorophore to the Aβ peptide.

### Preparation of HFIP-treated Aβ and Aβ^pH^ stocks

Aβ and Aβ^pH^ was dissolved in HFIP and prepared as previously described^34^. Briefly, 1 mM Aβ solution was prepared by adding HFIP directly to the vial (0.5 mg Aβ or Aβ^pH^ in 93.35 *μ*l HFIP). The peptide should be completely dissolved. The solution was incubated at room temperature for at least 30 min. HFIP was allowed to evaporate in the open tubes overnight in the fume hood and then dry down under high vacuum for 1 hour without heating to remove any remaining traces of HFIP and moisture. The resulting peptide thin clear film formed at the bottom of the tubes. The tubes containing dried peptides were stored at −20°C until further use. To make oligomers, 5 mM Aβ DMSO stock was prepared by adding 20 *μ*l fresh dry DMSO to 0.45 mg. To ensure complete resuspension of peptide film, this was pipetted thoroughly, scraping down the sides of the tube near the bottom. This was vortexed well (~30 s) and pulsed in a microcentrifuge to collect the solution at the bottom of the tube and the 5 mM Aβ DMSO solution was sonicated for 10 min. This preparation was used as the starting material for preparing the aggregated Aβ.

### Atomic Force Microscopy for analysis of Aβ and Aβ^pH^

We followed the previously published detailed protocols for analyzing Aβ and Aβ^pH^ by Atomic Force Microscopy (AFM)^34,56^. Oligomers were prepared by incubating the samples at 4°C for 24 hours. Briefly, sample preparation was performed with sterile techniques using sterile media and MilliQ- water. A 10 ml syringe with ultrapure water equipped with a 0.22 *μ*m filter was filled and the initial 1–2 ml was discarded though syringe filter output. 1M HCl and 1x PBS buffer were also filtered through 0.22 *μ*m filter. The samples were prepared for spotting on mica by diluting to final concentrations of 10–30 *μ*M in water. Immediately before sample delivery, top few layers of mica were cleaved away using an adhesive tape to reveal a clean, flat, featureless surface. The fresh surface was pretreated with ~5-8 *μ*l of filtered 1M HCl for 30 seconds and rinsed with two drops of water (note: the mica was held at a 45° angle and washed with water to allow the water coming out of the syringe filter to roll over the mica). If necessary, the remaining water was absorbed with fiber-free tissue paper/wipes by keeping paper on the edge of the mica. Immediately, the sample was spotted onto mica and incubated for 3 minutes followed by rinsing with three drops of water and blow drying with several gentle pulses of compressed air. Samples were then kept in a dust-free box and incubated on benchtop for a few minutes to hours at room temperature until analysis. AFM imaging was performed with Veeco Multimode with NanoScope V controller with NanoScope Software using the Silicon AFM probes, TAP300 Aluminum reflex coating (Ted Pella, Inc. Cat# TAP300AL-G-10) at ~300 kHz resonant frequency and ~40 N/m force constant in the tapping mode.

### pH-dependent emission spectra of Aβ conjugated with Protonex Green^®^ and Aβ conjugated with pHrodo at various concentrations

The cell culture medium was treated with dilute HCl and NaOH solutions to obtain different solutions of pH ranging from 1.0 to 9.0 for the assay. Lyophilized powder of Aβ conjugated with pHrodo (RODO-Aβ^pH^) and Aβ conjugated with Protonex Green^®^ (PTXG-Aβ^pH^) was dissolved in cell culture medium to make stock solutions and kept at 37°C for 24 hours to pre-aggregate the peptide conjugates. From the stock solutions, different dilutions for each pH condition was prepared at concentrations of 0.5, 1.0, 2.0, and 5.0 *μ*M in a 96-well plate (100 *μ*l/well). Fluorescence intensity of each well containing Aβ^pH^ was obtained on a Cytation™ 5 imaging multi-mode reader (BioTek Instruments) at 443/505 nm excitation/emission wavelengths. The fluorescence intensity of each pH-solution and Aβ^pH^-concentration in relative fluorescence units (RFU) was plotted using GraphPad Prism software.

### Concentration-dependent response of Protonex Green and Aβ^pH^ at acidic pH over time

The solutions of Protonex Green (PTXG) and Aβ conjugated with PTXG (Aβ^pH^) were prepared in the cell culture medium at concentrations of 0, 0.1, 0.5, 1.0, 2.0, 5.0 *μ*M and 50 *μ*l of each concentration were taken in duplicates in the wells of a 96-well plate. To measure the florescence of PTXG and Aβ^pH^ solutions in acidic conditions at different concentrations, 7.5 *μ*l of pH 1.0 solution (hydrochloric acid in media) was added to each well to obtain a final pH of 3.0. Fluorescence intensities of the acidic solutions were measured at an excitation/emission of 443/505 nm on A Cytation 5 multimode plate reader (BioTek, Inc). Next, to initiate aggregation of Aβ^pH^, the plate was incubated at 37°C, 5% CO_2_ and fluorescence was measured at 2, 6, 12, and 24-hour time points. The change in fluorescence intensities of the PTXG and Aβ^pH^ aggregates was analyzed over time using GraphPad Prism software.

### Cell lines – culture and maintenance

BV2 and N9 mouse microglial cell lines were generously gifted by Dr. Linda J. Van Eldik (University of Kentucky, USA). The BV-2 cell line was developed in the lab of Dr. Elisabetta Blasi at the University of Perugia, Italy. Cells were maintained at 37°C and 5% CO_2_ in DMEM (Dulbecco’s Modified Eagle’s Medium)/Hams F-12 50/50 Mix (Corning #10-090-CV) supplemented with 10% FBS (Atlanta Biologics), 1% L-Glutamine (Corning #25-005-CI), and 1% Penicillin/Streptomycin (Invitrogen). HMC3 human microglial cell line was a gift from Dr. Jianming Li (Purdue University, USA) who originally obtained the cells from ATCC. These cells were maintained at 37°C and 5% CO_2_ in DMEM supplemented with 10% FBS and 1% penicillin/streptomycin.

### Phagocytosis assay with live microglia

Cells were seeded at 5000 cells per well (200 *μ*l per well) in a 96-well flat bottom plate (Falcon) for approximately 16 hours (overnight). For all cell assays, the lyophilized Aβ^pH^ conjugate was dissolved in the culture medium to prepare a stock solution and was pre-aggregated by placing the stock solution at 37°C for 24 hours. Further dilution for cell treatment was performed in culture media and the diluted solution was filtered using 0.22 *μ*m syringe filter prior to cell treatment. The adherent cells (BV2, N9, HMC3) were treated with a final concentration of Aβ^pH^ at 0, 0.1, 0.5, 1.0, 2.0, and 5.0 *μ*M by replacing one-half of the culture medium (100 *μ*l) with 2x concentration. Two technical replicates were used for each treatment concentration. The plates were immediately placed in an IncuCyte S3 Live-Cell Analysis System (Essen BioScience) and four images per well were captured at 30-minute time intervals for 24 hours. The fluorescence intensity, cell confluence, and the integrated fluorescence intensity data was obtained and analyzed using the GraphPad Prism software.

The Aβ^pH^ uptake was measured as the *Phagocytic Score* metric, defined as a normalized value relative to the initial fluorescent intensity at t=0 and calculated as:

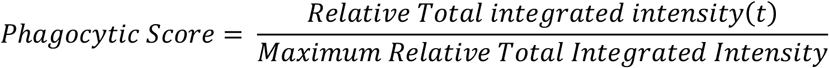

where, *Relative Total Integrated Intensity* is defined as *Total Integrated Intensity (t) – Total Integrated Intensity (t=0)* for each concentration and cell type.

The *Total integrated intensity* is defined as the total sum of Aβ^pH^ fluorescent intensity in the entire image and given by the expression 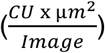 as defined by Incucyte. We captured 4 images per well with multiple replicates for each Aβ^pH^ concentration and for each cell type. These images were used to calculate the value of *Total integrated intensity*, 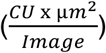. The individual units are defined as: CU = Average mean intensity (the average of Aβ^pH^’s mean fluorescent intensity in each cell in an image), μ*m*^2^= Average area (the average area of the Aβ^pH^’s in each cell in an image).

The *Maximum Relative Total Integrated Intensity* is the maximum value of *Relative Total Integrated Intensity* over the 24-hour period. Such a normalization gives *Phagocytic Score* values between 0 and 1 to compare different concentrations across different cell types. The maximum peak indicates the time when the degradation is equal to the uptake for each concentration and cell type. All other values show an interplay between uptake or degradation compared to time, t=0, shown by either increase or decrease in florescence that is also observed visually. The corresponding videos (**Movies 1-3**) during live cell imaging were taken on IncuCyte S3 Live-Cell Analysis System (Essen BioScience) and stabilized using the Blender version 2.82a software (www.blender.org).

### Isolation and culture of primary mouse microglia

CD11b^+^ primary microglia were isolated from adult mice aged around 7 months (male and female) and cultured as follows. Mice were euthanized with CO_2_ following the Purdue University Animal Care and Use Committee guidelines and brains were transcardially perfused with ice-cold PBS. The perfused brains were dissected and cut into small 1 mm^3^ pieces before homogenizing them in DPBS^++^ containing 0.4% DnaseI on the tissue dissociator (Miltenyi Biotec) at 37 °C for 35 mins. The cell suspension was filterered through a 70 *μ*m filter and myelin was removed two times, first using Percoll PLUS reagent followed by myelin removal beads using LS columns (Miltenyi Biotec). After complete myelin removal, CD11b^+^ cells were selected from the single cell suspension using the CD11b beads (Miltenyi Biotec) as per the manufacturer’s instructions. The CD11b^+^ cells were finally resuspended in microglia growth media made in DMEM/F12 (Corning Cat. #MT15090CV), further diluted in TIC (TGF-β, IL-34, and cholesterol) media^37^ with 2% FBS before seeding 0.1×10^6^ cells per 500 *μ*L in a well of a 24-well plate (Corning Cat. #353847). The cells were maintained in TIC media at 37 °C and 10% CO2 with media change every other day until the day (around 10-14 div) of the phagocytosis assay.

### Flow cytometry analysis of BV2 and primary microglia phagocytosis

BV2 microglial cells were seeded at a density of 250k cells/well in a 6-well plate for around 14 hours overnight. The next morning, the cells were treated with 0.5 *μ*M and 5 *μ*M Aβ^pH^ and placed in a 37°C incubator for 1 hour, after which the plate was brought to the hood, placed on ice to stop phagocytosis, and cell culture medium containing Aβ^pH^ was aspirated. The cells were washed once with cold PBS. Next, the cells were treated with ice cold PBS containing 2 mM EDTA for 2 mins on ice to detach the cells from the wells. The cells were then centrifuged at 1400 rpm for 3 mins. The supernatant was aspirated, and the cell pellets were re-suspended in FACS buffer (PBS, 25 mM HEPES, 2mM EDTA, and 2% FBS). Five minutes before analysis of each sample, propidium iodide (PI; Thermo Fisher Scientific, Cat. #P1304MP) was added to the sample (40 ng/ml cell suspension) for staining of dead cells.

Phagocytosis of Aβ^pH^ by primary cells was performed on div 10-14 in a similar manner. Cells were treated with Aβ^pH^ for 1 hour and detached from the plate using cold PBS and 2 mM EDTA. After centrifuging the cell suspension, the cell pellet was resuspended in 0.1 mL PBA for live/dead staining. Here, Zombie Violet, ZV, (BioLegend, #423113) was used to evaluate cell viability (1:100 per 10^6^ cells in 0.1 mL) for 15 mins followed by PBS wash. Finally, the cells were resuspended in FACS buffer and taken for analysis. Cells exhibiting green fluorescence were captured on the FITC channel upon gating for live cells (PI^-^ negative or ZV^-^ cells are taken as live cells) on Attune NxT flow cytometer (Invitrogen). The files were then analyzed on FlowJo V10 software. Briefly, at least 90% of the total cells were first gated on SSC-A vs FSC-A plot and the single cells within this gate were selected on the FSC-H vs FSC-A plot. From the single cells, the live and dead cells were identified from the viability dye. PI- and ZV- cells were considered as live cells and PI^+^ and ZV^+^ cells were taken as dead cells on the histogram plots. Finally, Aβ^pH+^ or Aβ^pH-^ cells were identified within the live cell population in the Aβ^pH^ histogram plot on the FITC channel.

### Confocal imaging of actin filaments and nuclei in the paraformaldehyde-fixed phagocytic microglial cells

For labeling the cells with phalloidin and DAPI, 20,000 cells/250 *μ*l were plated in 14 mm microwells of 35 mm glass bottom dishes (MatTek #P35G-1.5-14-C) and kept overnight. The cells were treated with 5.0 *μ*M Aβ^pH^ for 2 hours on the next day. Then the medium was aspirated, and cells were fixed with 4% paraformaldehyde for 20 mins, and then gently washed once with PBS. Phalloidin-iFluor 594 reagent (Abcam, Cat. #ab176757) was diluted at a concentration of 1 *μ*l/ml in PBS and added to the fixed cells for 10 minutes for staining the actin filaments. To label the nuclei, DAPI was diluted at a concentration of 1 *μ*l/ml. The cells were washed again with PBS followed by a 10-minute incubation with the diluted DAPI solution. Finally, the DAPI solution (Invitrogen, Cat. #D3571) was aspirated and the fixed cells treated with 2-3 drops of ProLong Gold Antifade Mountant (Invitrogen #P36930) before imaging. Fluorescence images of phagocytic microglial cells were captured using 40x and 60x objectives on the Nikon AR-1 MP confocal laser microscope. Images were obtained on the NIS Elements microscope imaging software.

### Confocal Imaging of intracellular acidic organelles and nuclei in paraformaldehyde-fixed phagocytic microglial cells

LysoTracker Red DND-99 (Thermo Fisher Scientific, Cat. #L7528) was used for labeling the intracellular organelles of the cells to observe localization of Aβ^pH^ sensors inside the cells after phagocytosis. 20,000 cells/250 *μ*l were plated in 14 mm microwells of 35 mm glass bottom dishes (MatTek #P35G-1.5-14-C) and kept overnight. The cells were treated with 5.0 *μ*M Aβ^pH^ for 2 hours on the next day. Then the Aβ^pH^ containing medium was aspirated and replaced with 200 *μ*l of media containing 100 nM concentration of the LysoTracker dye and the cells were incubated in the 37°C, 5% CO_2_ incubator for 30 minutes. Finally, the cells were fixed and treated with DAPI to stain the nuclei followed by 2-3 drops of ProLong Gold Antifade Mountant using the above-mentioned protocol. Fluorescence images of phagocytic microglial cells were captured using 40x and 60x objectives on the Nikon AR-1 MP confocal laser microscope. Images were obtained on the NIS Elements microscope imaging software.

### Intracranial Injection of Aβ^pH^, perfusion and immunohistochemistry

Lyophilized Aβ^pH^ was dissolved in Hank’s Balanced Salt Solution (HBSS no calcium, no magnesium, no phenol red, ThermoFisher #14175079) to obtain a 100 *μ*M stock solution that was then briefly vortexed and sonicated in a bath sonicator for 1 minute and directly used for intracranial injections or stored at −80°C. For intracranial injections, the stock solution was diluted in HBSS to obtain a 10 *μ*M working solution and kept on ice to prevent aggregation. Postnatal day 7 mice were anesthetized with isoflurane and were mounted on a stereotactic frame. Two lots of 250 nl of Aβ^pH^ working solution were unilaterally injected in the somatosensory cortex of wild-type C57BL/6J mice using a Nanoliter Injector (anteroposterior −2.3 mm; mediolateral +2.3 mm; dorsoventral −0.5 mm for lower layers and −0.2 mm for upper layers, relative to Lambda) at an injection rate of 100 nl/minute followed by 2 additional minutes to allow diffusion. 24 hours or 72 hours after injections, animals were deeply anesthetized with sodium pentobarbital by intraperitoneal injection and then transcardially perfused with PBS 1x followed by 4% paraformaldehyde (PFA) in PBS. Brains were dissected out and post-fixed for two hours at 4°C, cryoprotected in a 30% sucrose-PBS solution overnight at 4°C. Then, tissue was sectioned at 40 *μ*m on a sliding microtome (Leica). Free-floating brain sections were permeabilized by incubating with 0.3% Triton X-100 in PBS for 1 hour and then blocked for 3 hours (0.3% Triton X-100 and 10% normal donkey serum), followed by incubation with primary antibodies in 0.3% Triton X-100 and 10% normal donkey serum overnight at 4°C. The next day, brains were rinsed in PBS 1x for 1 hour, incubated with the appropriate secondary antibodies for 2 hours at room temperature, rinsed again in PBS, incubated with DAPI and mounted using Fluoromount-G (SouthernBiotech, #0100-01). During perfusion and immunohistochemistry all solutions were maintained at neutral pH. The following primary antibodies were used: mouse anti-GFAP (1:200, Sigma #G3893-100UL) and rabbit anti-IBA1 (1:500, Wako Chemicals, #019-1974). The secondary antibodies used were donkey anti-mouse-IgG1 647 (Invitrogen, #A21241) and donkey anti-rabbit 594 (ThermoFisher, #A-21207). Tissue samples were imaged on a ZEISS Axio Imager and a ZEISS LSM 800 confocal using a 20X objective. Aβ^pH^ fluorescence signal was quantified using ImageJ. First, cell contour was manually drawn using IBA1 or GFAP signal and mean fluorescence intensity in the Aβ^pH^ channel was measured. Next, the selection was moved to a nearby region with no obvious Aβ^pH^ fluorescence signal and mean fluorescence intensity in the Aβ^pH^ channel was measured (background). Normalized fluorescence intensity for each cell was calculated as Aβ^pH^ fluorescence signal minus background. Data were analyzed by one-way ANOVA followed by the Sidak’s post hoc analysis for comparisons of multiple samples using GraphPad Prism 7 (GraphPad Software).

### Immunopanning and culture of primary astrocytes

Astrocytes were purified by immunopanning from P5 Sprague Dawley rat (Charles River) forebrains and cultured as previously described^57^. In brief, cortices were enzymatically (using papain) then mechanically dissociated to produce a single-cell suspension that was incubated on several negative immunopanning plates to remove microglia, endothelial cells and oligodendrocyte lineage cells. Positive selection for astrocytes was with an ITGB5-coated panning plate. Isolated astrocytes were cultured in a defined, serum-free base medium containing 50% neurobasal, 50% DMEM, 100 U ml^−1^ penicillin, 100 *μ* g ml^−1^ streptomycin, 1 mM sodium pyruvate, 292 *μ*g/ml L-glutamine, 1× SATO and 5 *μ*g/ml of *N*-acetyl cysteine. This medium was supplemented with the astrocyte-required survival factor HBEGF (Peprotech, 100-47) at 5 ng/ml^57^. Cells were plated at 5,000 cells/well in 12 well plates coated with poly-D-lysine and maintained at 10% CO_2_.

### Engulfment assay of Aβ^pH^ by primary astrocytes

Astrocytes were maintained for 1 week in culture, checked for reactivity state using qPCR^5^ before addition of Aβ^pH^. The cells were treated with 0.5, 1.0, or 2.0 *μ*M Aβ^pH^ and imaged continuously with still images taken every 5 minutes with an IncuCyte S3 System epifluorescence time lapse microscope to analyze engulfed Aβ^pH^ particles. For image processing analysis, we took 9 images per well using a 20× objective lens from random areas of the 12-well plates and calculated the phagocytic index by measuring the area of engulfed synaptosomes/myelin (fluorescent signal) normalized to the area of astrocytes, using ImageJ.

### *In vivo* retinal engulfment of Aβ^pH^

P14 Sprague Dawley rats were anaesthetized with 2.5% inhaled isoflurane in 2.0 l O2 per min. Once non-responsive, animals received a 1 *μ*l intravitreal injection of Aβ^pH^, or PBS. Retinae were collected for immunofluorescent analyses at 3, 24, and 72 hours. At collection, eyeballs were removed, fixed in 4% PFA overnight, washed in PBS and retinae dissected and whole-mounts placed on silanized glass slides. Retinae were blocked with 10% heat-inactivated normal goat serum for 2 hours at room temperature. Incubation with primary antibodies to GFAP (Dako, A0063, 1:5000) and IBA1 (WAKO, 019-19741, 1:500) diluted in 5% goat serum in PBS was followed by detection with AlexaFluor fluorescent secondary antibodies (Thermo, 1:1000).

### Brain slice experiments

Rats at postnatal day 12 (P12) were sacrificed by cervical dislocation followed by decapitation, and 250 *μ*m sagittal hippocampal slices were prepared on a Leica VT 1200S vibratome at 4ºC in oxygenated solution containing (mM): 124 NaCl, 26 NaHCO_3_, 2.5 KCl, 1 NaH_2_PO_4_, 10 glucose, 2 CaCl_2_, 1 MgCl_2_, 1 kynurenic acid. Acute slices were allowed to recover for 2.5 hours at room temperature before being transferred to 24-well plates and incubated with 5 *μ*M Aβ^pH^ in HEPES-based aCSF (140 mM NaCl, 10 mM HEPES, 2.5 mM KCl, 1 mM NaH_2_PO_4_, 10 mM glucose, 2 mM CaCl_2_, and 1 mM MgCl_2_) for 1.5 hours at 37 ºC. Following incubation, slices were quickly rinsed in cold phosphate-buffered saline (PBS) and fixed in 4% paraformaldehyde (PFA) for 45 min at room temperature. For immunolabeling, slices were permeabilized and blocked in buffer containing 10% horse serum and 0.02% Triton X 100 in PBS for 2 hours at room temperature, followed by incubation with goat anti-IBA1 (Abcam, ab5076) or chicken anti-GFAP (Abcam, ab4674) primary antibodies diluted 1:500 in blocking buffer for 12 h at 4°C. Following four 10 min washes in PBS, donkey anti-goat IgG 647 (ThermoFisher, A21447) or donkey anti-chicken IgG 649 (Jackson, 703-495-155) secondary antibodies diluted 1:1000 in blocking buffer were applied for 4 h at room temperature. Finally, slices were incubated in DAPI for 30 min, rinsed in PBS and mounted. Imaging was done using a Zeiss LSM700 confocal microscope and a 20X objective, where 10 *μ*m image stacks at 1 *μ*m step interval were acquired. For colocalization analysis, the percentage of Aβ^pH^ signal within microglial or astroglial cells was calculated by binarizing the microglia or astrocyte channel, creating a mask and multiplying it by the raw Aβ^pH^ signal. The fluorescence intensity of Aβ^pH^ colocalizing with either cell type mask was then expressed as a percentage of the total Aβ^pH^ signal across the field.

### *In vivo* two-photon microscopy

Adult mice bred on a C57BL/6 background aged ~4-6 months (P123 to P203) were anesthetized using urethane (1.55 g/kg given intraperitoneally). Adequate anesthesia was ensured by confirming the absence of a withdrawal response to a paw pinch. Body temperature was maintained at 36.8±0.3°C and eyes were protected from drying by applying polyacrylic acid eye drops (Dr Winzer Pharma). The trachea was cannulated and mice were mechanically ventilated with medical air supplemented with oxygen using a MiniVent (Model 845). A headplate was attached to the skull using superglue and mice were head fixed to a custom-built stage. A craniotomy of approximately 2 mm diameter was performed over the right primary somatosensory cortex, immediately caudal to the coronal suture and approximately 2 to 4 mm laterally from the midline. The dura was removed and 2% agarose in HEPES-buffered aCSF was used to create a well filled with HEPES-buffered aCSF during imaging.

Two-photon excitation was performed using a Newport-Spectraphysics Ti:sapphire MaiTai laser pulsing at 80 MHz, and a Zeiss LSM710 microscope with a 20× water immersion objective (NA 1.0). Fluorescence was evoked using a wavelength of 920 nm. The mean laser power under the objective did not exceed 25 mW. Image stacks were taken in 2 *μ*m depth increments (50-200 *μ*m from the cortical surface) every 1.5 min for approximately 30 min. Five *μ*M Aβ^pH^ in HEPES-based aCSF was applied to the cortical surface for 10 min and then replaced with HEPES-based aCSF. Animals were transcardially perfused with ice-cold PBS followed by 4% PFA in PBS at 1.5 or 3 h after pH Aβ^pH^ application. Brains were post-fixed in PFA for 12 h at 4°C and 100 *μ*m sagittal sections were prepared using a vibratome. Slices were permeabilized and blocked for 12 h and incubated with rabbit anti-IBA1 (1:500, Synaptic Systems, 234006), rat anti-mouse CD68 (1:500, Bio-Rad, MCA1957) or chicken anti-GFAP (1:500, Abcam, ab4674) primary antibodies for 24 hours at 4°C. Following washes in PBS, donkey anti-rabbit IgG 647 (1:500, ThermoFisher, A31573), goat anti-rat IgG 647 (1:500, ThermoFisher, A21247) or donkey anti-chicken 649 IgG (1:300, Jackson, 703-495-155) were applied for 12 hours at 4°C. Image stacks (23 *μ*m deep) were acquired at 1 *μ*m interval in the cerebral cortex and analysis was done as for *in situ* experiments.

## Associated content

Supporting Information: Instrumentation used for Chemical Characterization, Supporting Movies and Imaging: live cell images, confocal images, videos of phagocytosis in real time, synthetic procedures and detailed characterization of Aβ^pH^, etc. are available in the Supporting Information file.

## Acknowledgment

This work was supported, in part, by a Purdue University start-up package from the Department of Chemistry at Purdue University, an award from Purdue Research Foundation, Ralph W. and Grace M. Showalter Research Trust award, the Jim and Diann Robbers Grant for New Investigators award and National Institutes of Health, National Center for Advancing Translational Sciences ASPIRE Design Challenge awards to Gaurav Chopra. Additional support was, in part by, the Stark Neurosciences Research Institute, the Indiana Alzheimer Disease Center, Eli Lilly and Company, the Indiana Clinical and Translational Sciences Institute grant # UL1TR002529 from the National Institutes of Health, National Center for Advancing Translational Sciences, and the Purdue University Center for Cancer Research funded by National Institutes of Health grant # P30 CA023168 are also acknowledged. The content is solely the responsibility of the authors and does not necessarily represent the official views of the National Institutes of Health. Work in the Attwell lab was supported by Wellcome Trust Investigator Awards 099222 and 219366 to David Attwell, by a Wellcome Trust 4-year PhD studentship to Pablo Izquierdo, and by a BBSRC LIDo PhD studentship to Nils Korte. Emilia Favuzzi was supported by EMBO (ALTF 444-2018). Kevin A. Guttenplan was supported by the Wu Tsai Neuroscience Institute Interdisciplinary Scholar Award from Stanford University. Work in the Liddelow lab was supported by NYU School of Medicine, generous anonymous donors, the Blas Frangione Foundation, and the Cure Alzheimer’s Foundation. We also thank Dr. Andy Schaber, Imaging Facility Director, Bindley Bioscience Center at Purdue University and Dr. Joydeb Majumder for assistance with confocal imaging; Prof. J. Paul Robinson and Kathy Ragheb at the Purdue University Cytometry Laboratories for flow cytometry assistance. The cartoon schematics in the figures were created using BioRender.

