## Supporting Information for "Monitoring phagocytic uptake of amyloid β into glial cell lysosomes in real time"

### INSTRUMENTATION USED FOR CHEMICAL CHARACTERIZATION OF Aβ$^{pH}$

1. MALDI spectra were obtained using Applied Biosystems Voyager DE PRO instrument (Main parameters: Number of laser shots: 100/spectrum, Laser intensity: 2977, Laser repetition rate: 20.0 Hz, accelerating voltage 25000 V).
2. $^{1}$H-NMR spectra were obtained in DMSO-$d_6$ solvent using a Bruker AV-III-500-HD 500 MHz NMR instrument.
3. ATR-FTIR spectra were recorded using Thermo Fisher Nicolet FTIR instrument.
4. AFM images were recorded using Veeco Multimode instrument with NanoScope V controller

### SUPPORTING MOVIES

**Movie S1: Phagocytosis of Aβ$^{pH}$ by HMC3 cells**. Phagocytic uptake of Aβ$^{pH}$ by HMC3 cells. The cells internalize Aβ$^{pH}$ and show increased green fluorescence with uptake over time.

**Movie S2: Phagocytosis of Aβ$^{pH}$ by BV2 cells.** Phagocytic uptake of Aβ$^{pH}$ by BV2 microglia is confirmed by the appearance of green fluorescence within the acidic phagosomes of the cells. A change from the ramified state to the amoeboid state is observed visually during this process.

**Movie S3: Phagocytosis of Aβ$^{pH}$ by N9 cells.** Phagocytic uptake of Aβ$^{pH}$ by N9 microglia demonstrated by the appearance of green fluorescence within the cells.

**Movie S4: *In vivo* imaging of Aβ$^{pH}$ in mouse cortex.** *In vivo* two-photon imaging of the barrel cortex before and after topical application of Aβ$^{pH}$ for 16 frames (intensity enhanced).

**Movie S5: *In vivo* imaging of Aβ$^{pH}$ in mouse cortex.** *In vivo* two-photon imaging of the barrel cortex before and after topical application of Aβ$^{pH}$. The fluorescence increase in cell somata (indicated by red circles) indicates Aβ$^{pH}$ uptake.

### SUPPORTING TEXT, IMAGES AND FIGURE LEGENDS

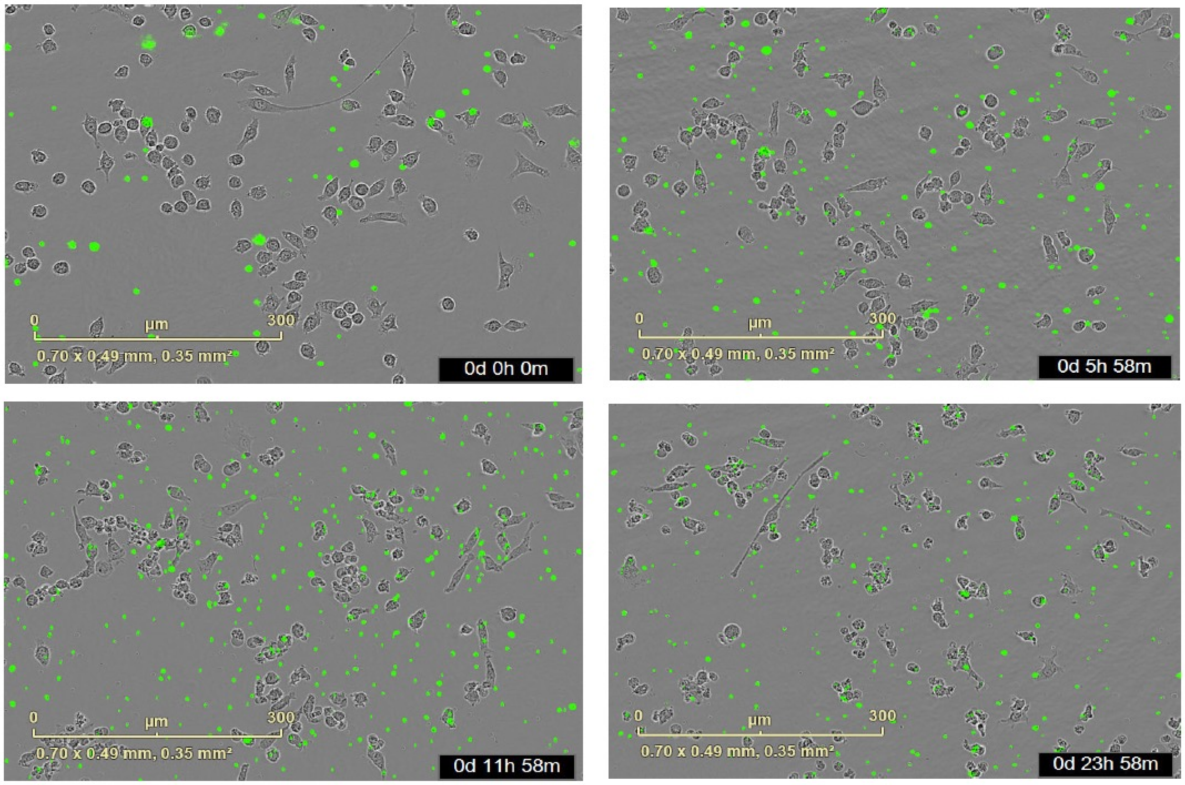


**Figure S1(a).** Live cell imaging of 1 µm fluorescent latex beads (Sigma-Aldrich, #L1030) applied to BV2 cells for 24 hours (~ 0, 6, 12, and 24 h). It is difficult to determine if the beads are within or outside the cell bodies.

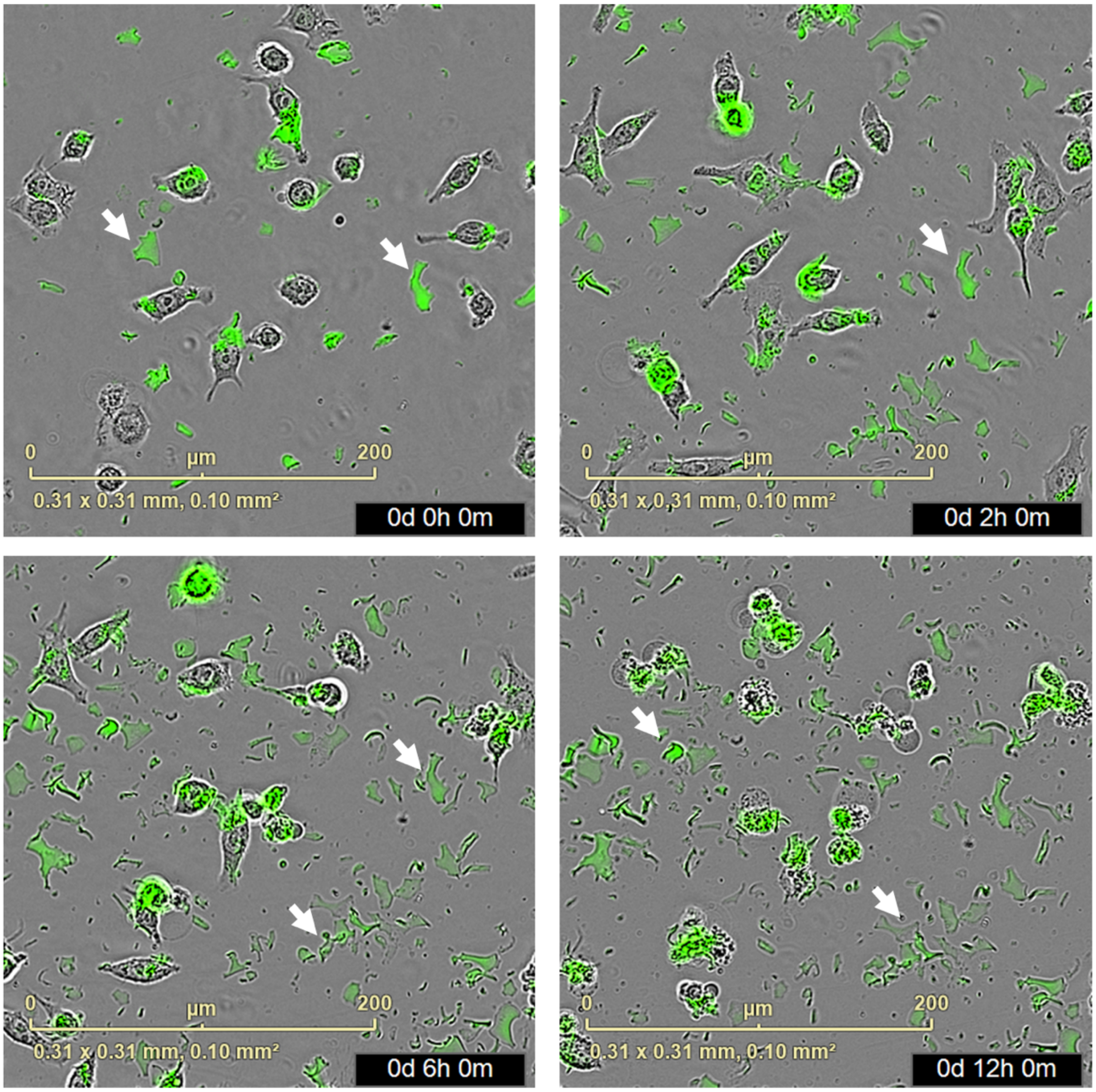


**Figure S1(b).** Live cell imaging of BV2 mouse microglia treated with (pH-independent) fluorescein-labeled Aβ peptides (AnaSpec, Inc. #23525-05) at 0, 2, 6, and 12 h time points. High background noise is observed in addition to non-specific fluorescence in and around the microglial cells.

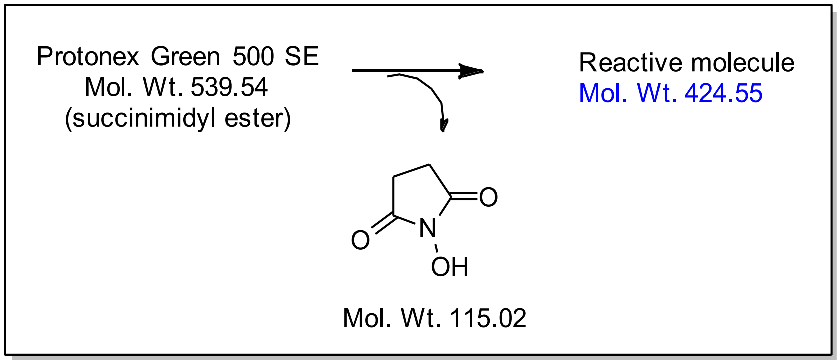


**Figure S2.** The mass calculation for the reactive part of Protonex Green 500, SE after conjugation with an amine functional group of Aβ$_{1-42}$. The Protonex Green 500, SE is a succinimidyl ester of active dye molecule (molecular weight 539.54) and the leaving group *N*-hydroxy succinimide has a molecular weight of 115.02. Therefore, the reactive fragment has a molecular weight of 424.55.

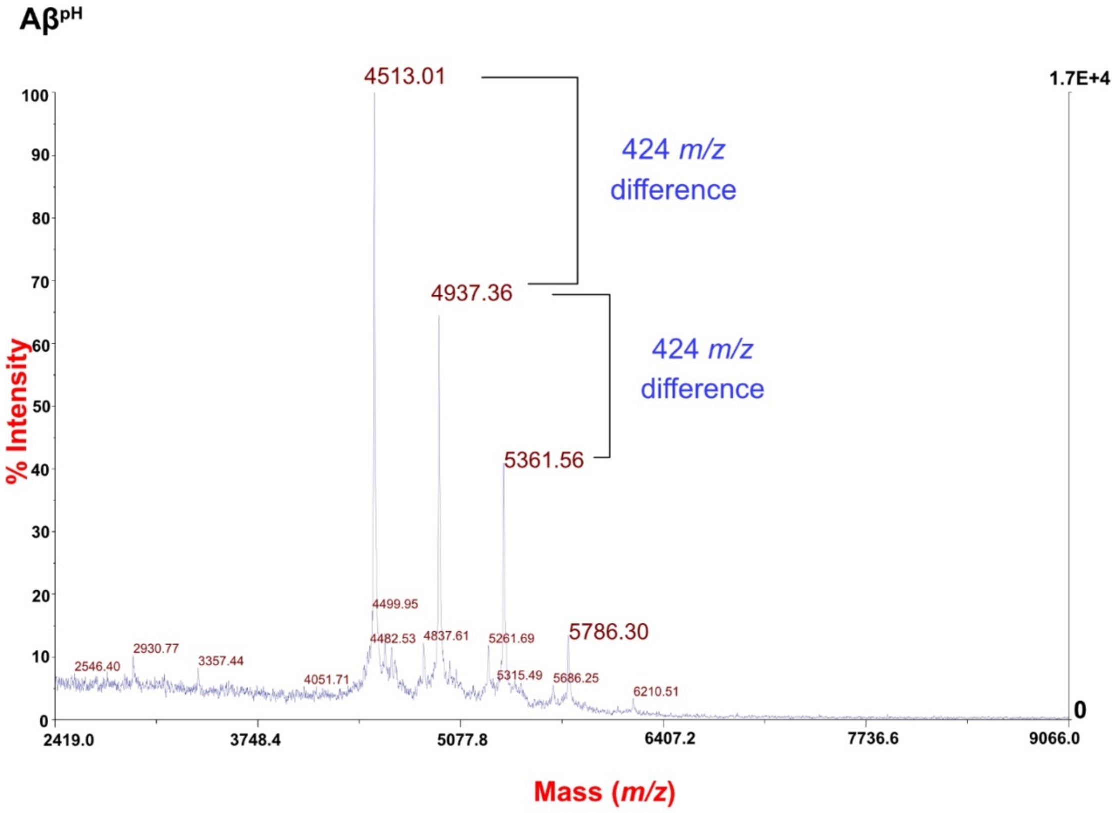


**Figure S3.** MALDI-MS spectrum [%intensity *vs.* Mass (*m/z*)] for Aβ-Protonex Green conjugate (Aβ$^{pH}$) corresponding to *m/z* of 5357.06 and 4932.87. The *m/z* difference of 424 indicates the addition of reactive fragment of Protonex Green 500, SE upon conjugation with an amine functional groups present in Aβ$_{1-42}$.

Proton nuclear magnetic resonance ($^{1}$H-NMR) spectrum shows a chemical shift for protons. A chemical shift is a relative resonant frequency to a standard magnetic field. Chemical shift (*δ*) is usually expressed in parts per million (ppm). The position and number of chemical shifts are used to determine the structure and functional groups present in a molecule. For example, chemical shifts for aliphatic protons are in 0.5 to 5.0 ppm region and for aromatic protons are in 6.5 to 9 ppm region depending upon the electrochemical environment of protons. This basis was used to identify aliphatic and aromatic protons in the $^{1}$H-NMR spectrum of PTXG, Aβ$_{1-42}$, PTXG-Aβ$^{pH}$ shown in **Figures S4-S6**, respectively.

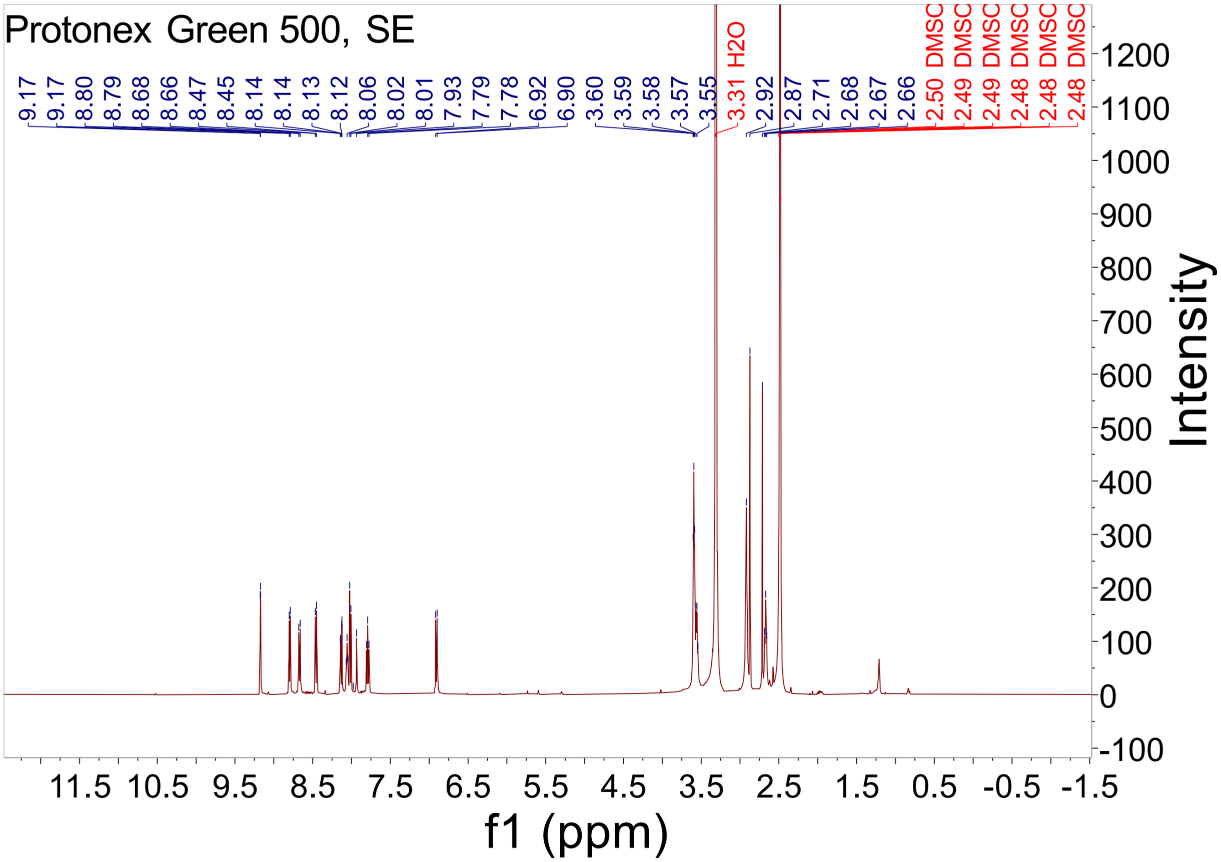


**Figure S4.** $^{1}$H-NMR spectrum [Intensity *vs.* ppm] of the Protonex Green 500, SE in DMSO-$d_6$. shows the presence of aliphatic (2.0 to 4.0 ppm region) and aromatic protons (6.5 to 9.5 ppm region to chemically characterize the material used for the reaction.

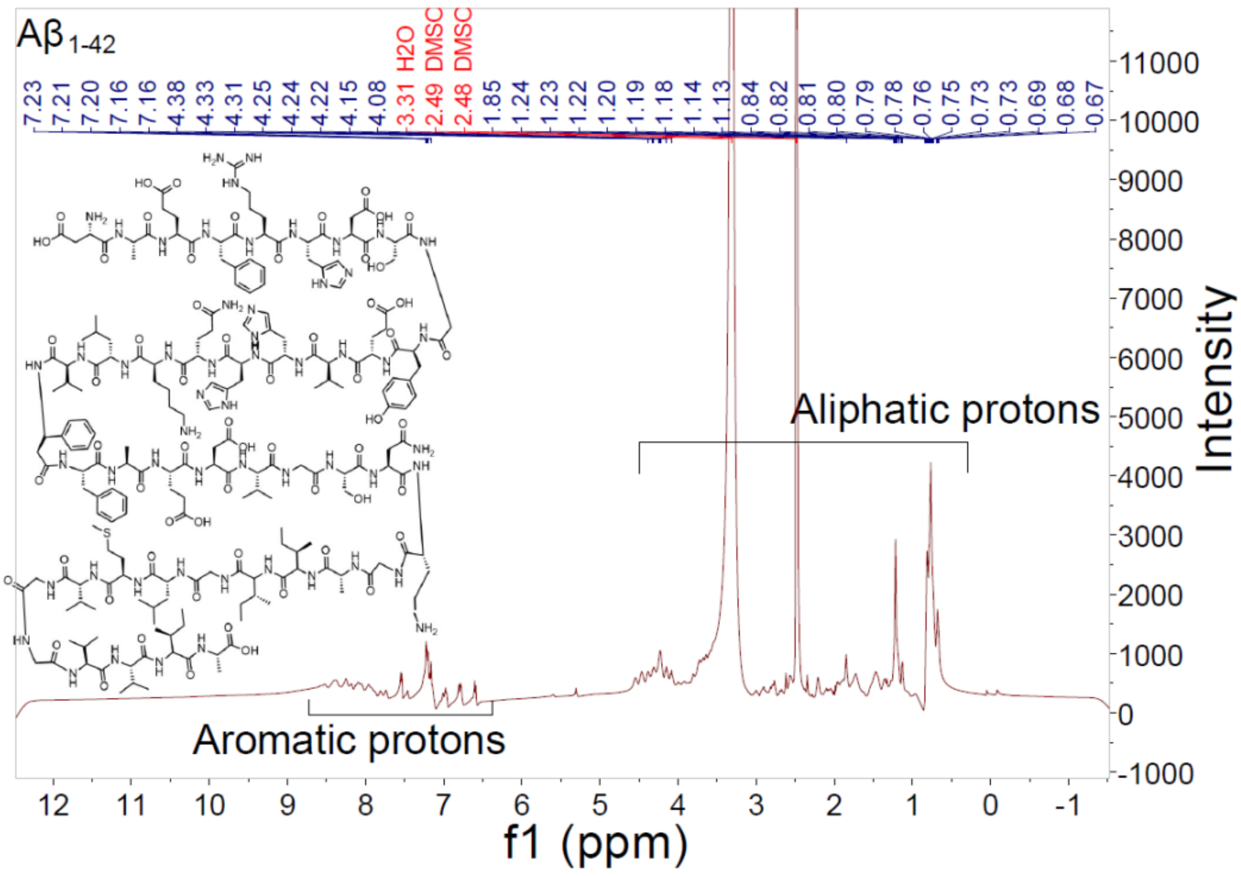


**Figure S5.** $^1$H-NMR spectrum [Intensity *vs.* ppm] of A$\beta_{1\text{-}42}$ in DMSO-$d_6$. The A$\beta_{1\text{-}42}$ peptide has amino acids containing aromatic (phenyl, 4-hydroxyphenyl, imidazolyl rings) and aliphatic (side chains and peptide backbone) functional groups. The $^1$H-NMR spectrum of A$\beta_{1\text{-}42}$ confirms the presence of protons arising from aliphatic and aromatic functional group-containing amino acids to characterize the material used for the reaction.

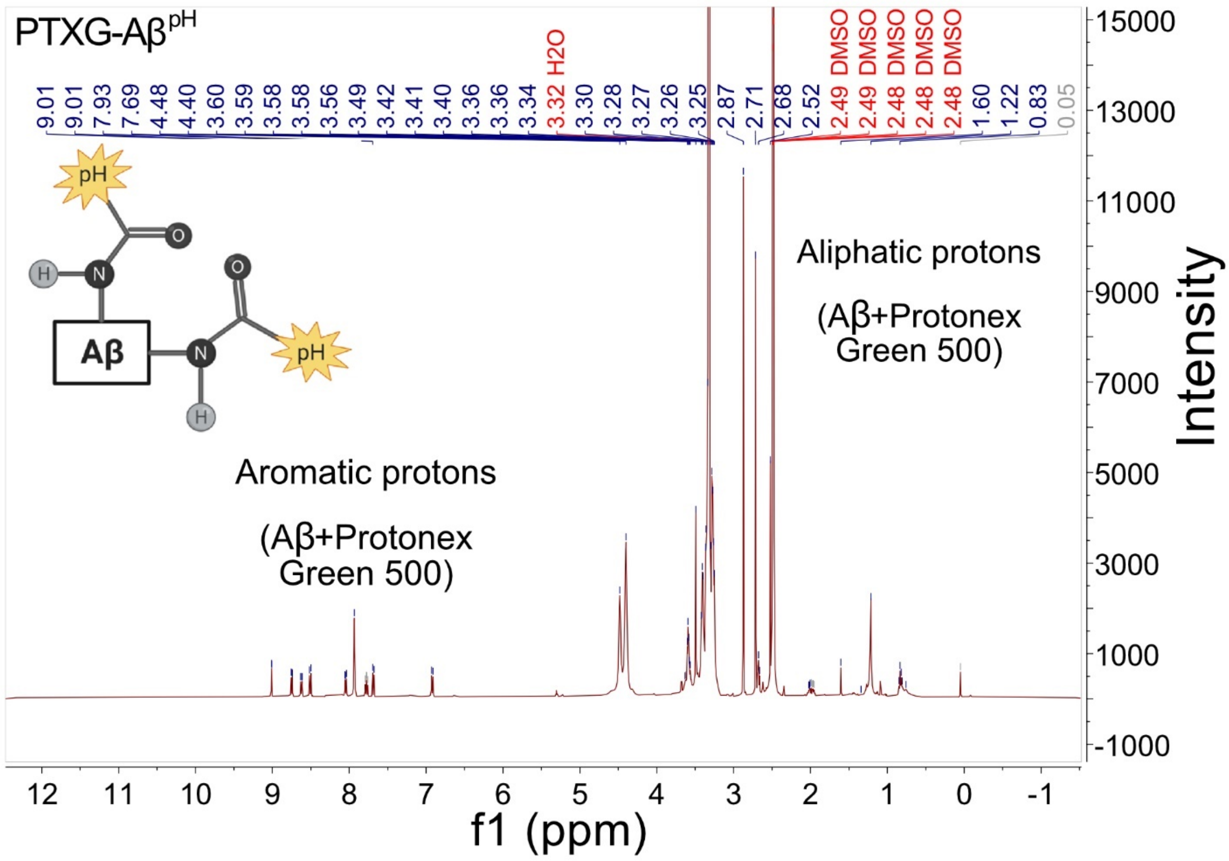


**Figure S6.** [1]H-NMR spectrum [Intensity *vs.* ppm] of the synthetic Protonex Green-conjugated Aβ (PTXG-$A\beta^{pH}$) in DMSO-$d_6$. The spectrum exhibits peaks at aliphatic and aromatic regions that are present due to functional groups present in both Protonex Green 500 and $A\beta_{1-42}$ (compare to **Figures S4, S5**) that confirms the conjugation of Protonex Green 500 with $A\beta_{1-42}$.

Fourier Transform InfraRed (FTIR) spectroscopy utilizes the frequencies associated with the bonds in a molecule that typically vibrate around 4000 $cm^{-1}$ to 400 $cm^{-1}$, known as the Infrared region of the electromagnetic spectrum. This region is associated with specific frequencies that change the vibration patterns of chemical bonds, resulting in an FTIR spectrum. A typical FTIR spectrum is visualized in a graph of infrared light absorbance (or transmittance) on the y-axis *vs.* frequency or wavenumber ($cm^{-1}$) on the x-axis. In general, the FTIR-spectrum is unique for an individual chemical entity and a change in structure or a functional group can be identified by comparing changes in these spectra. For example, carbonyl (C=O) is a functional group that is easily identified by FTIR spectroscopy due to a prominent bond stretching vibration peak at unique wavenumber range of around 1670-1820 $cm^{-1}$. Since carbonyl (C=O) groups are present on carboxylic acids, aldehydes, amides, anhydrides, esters, and ketones, unique wavenumbers in FTIR spectrum confirm these specific types of carbonyl groups. Furthermore, chemical modification results in a change in wavenumber to identify changes between specific carbonyl group types. This basis was used to identify chemical changes in the Attenuated Total Reflection Fourier Transform InfraRed (ATR-FTIR) spectrum of $A\beta_{1-42}$, PTXG, PTXG-$A\beta^{pH}$ shown in **Figures S7-S8**, respectively.

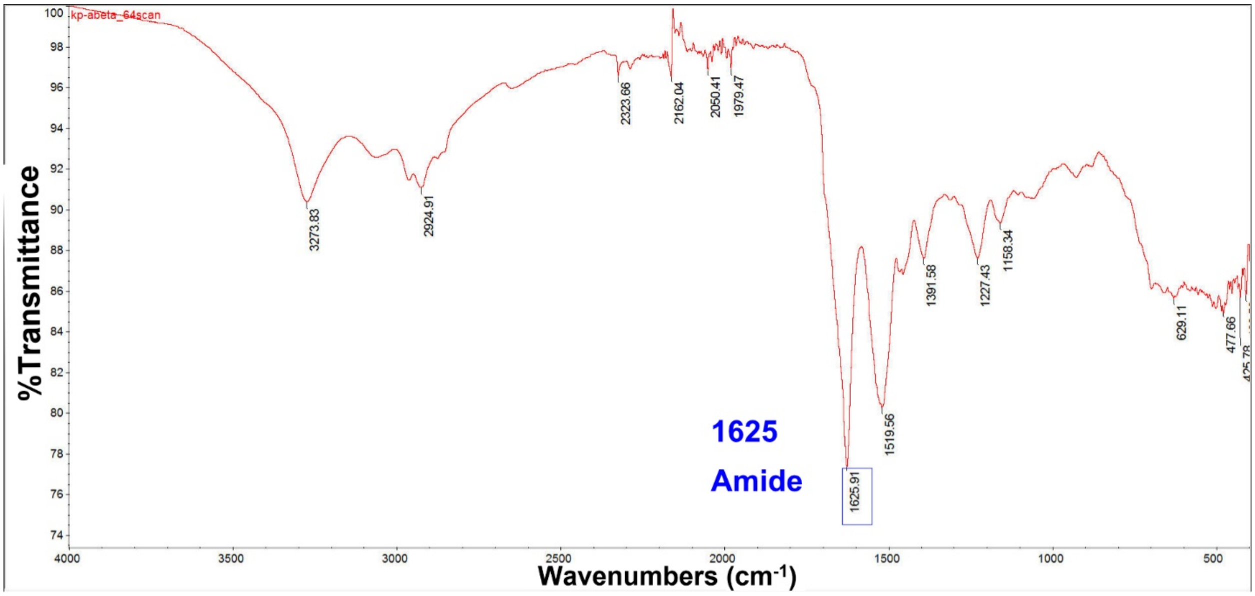


**Figure S7.** ATR-FTIR spectrum [%Transmittance *vs.* Wavenumbers ($cm^{-1}$)] of the $A\beta_{1-42}$ The ATR-FTIR shows the intense signal at 1625 $cm^{-1}$ correspondence to carbonyl (C=O) stretching frequencies in amide bonds of the peptide.

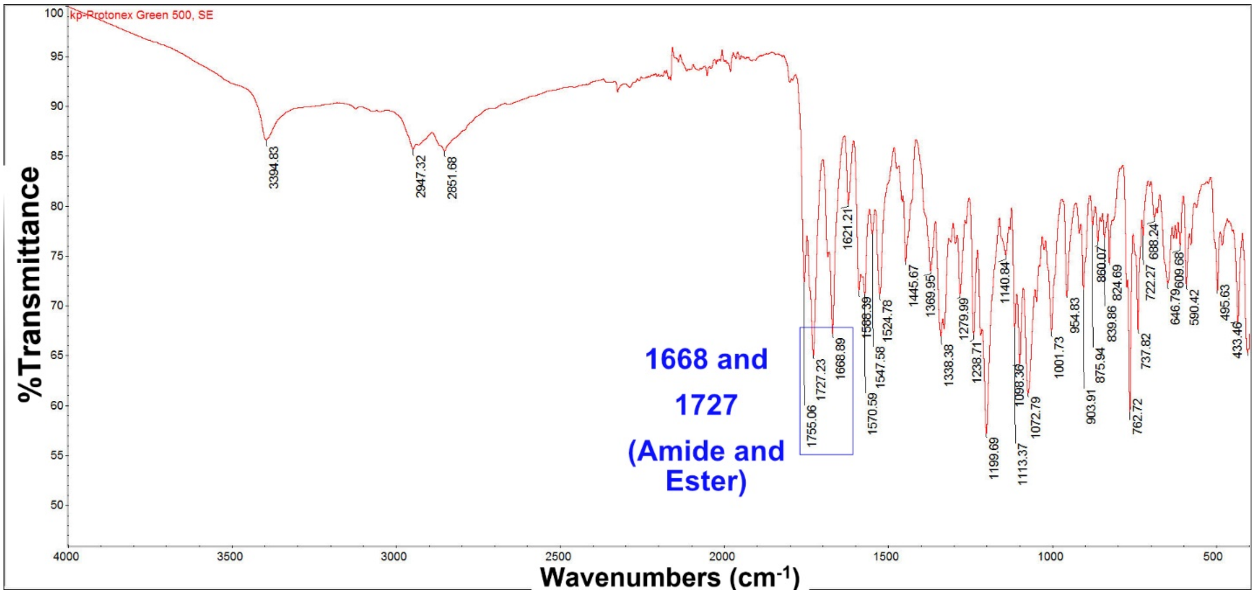


**Figure S8.** ATR-FTIR spectrum [%Transmittance *vs.* Wavenumbers ($cm^{-1}$)] of the Protonex Green 500, SE (PTXG). The PTXG chemical structure has amide and ester functional groups and the spectrum shows expected two intense signals at 1668 and 1727 $cm^{-1}$ corresponding to carbonyl (C=O) stretching frequencies in amide and ester functional groups, respectively.

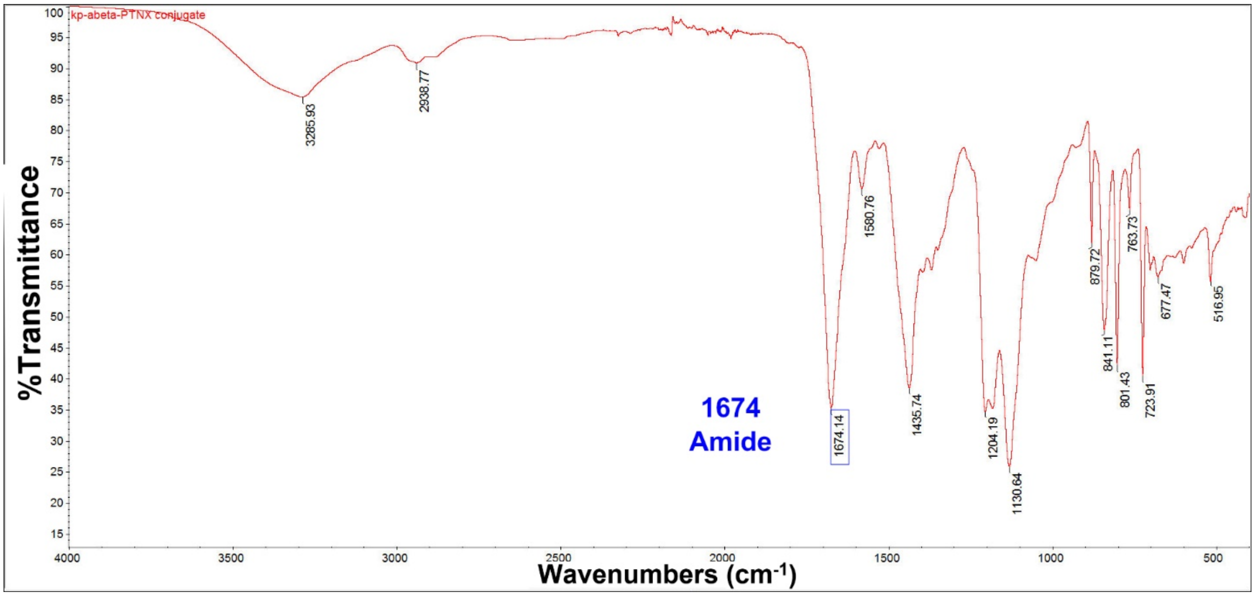


**Figure S9.** ATR-FTIR spectrum [%Transmittance *vs.* Wavenumbers ($cm^{-1}$)] of the PTXG-Aβ conjugate (PTXG-$A\beta^{pH}$) shows the change in stretching frequencies of carbonyl (C=O) of the amide functional group compared to **Figures S7-S8**. No peak at 1727 $cm^{-1}$ of the ester functional group as shown in **Figure S8**, suggests a change in the chemical structure of PTXG because of amide bond formation with an amine functional group of $A\beta_{1-42}$.

Taken together **Figures S4-S9** show chemical characterization of unconjugated Aβ, PTXG and conjugated PTXG-Aβ using $^1$H-NMR and ATR-FTIR spectroscopy.

**RODO-Aβ^pH**

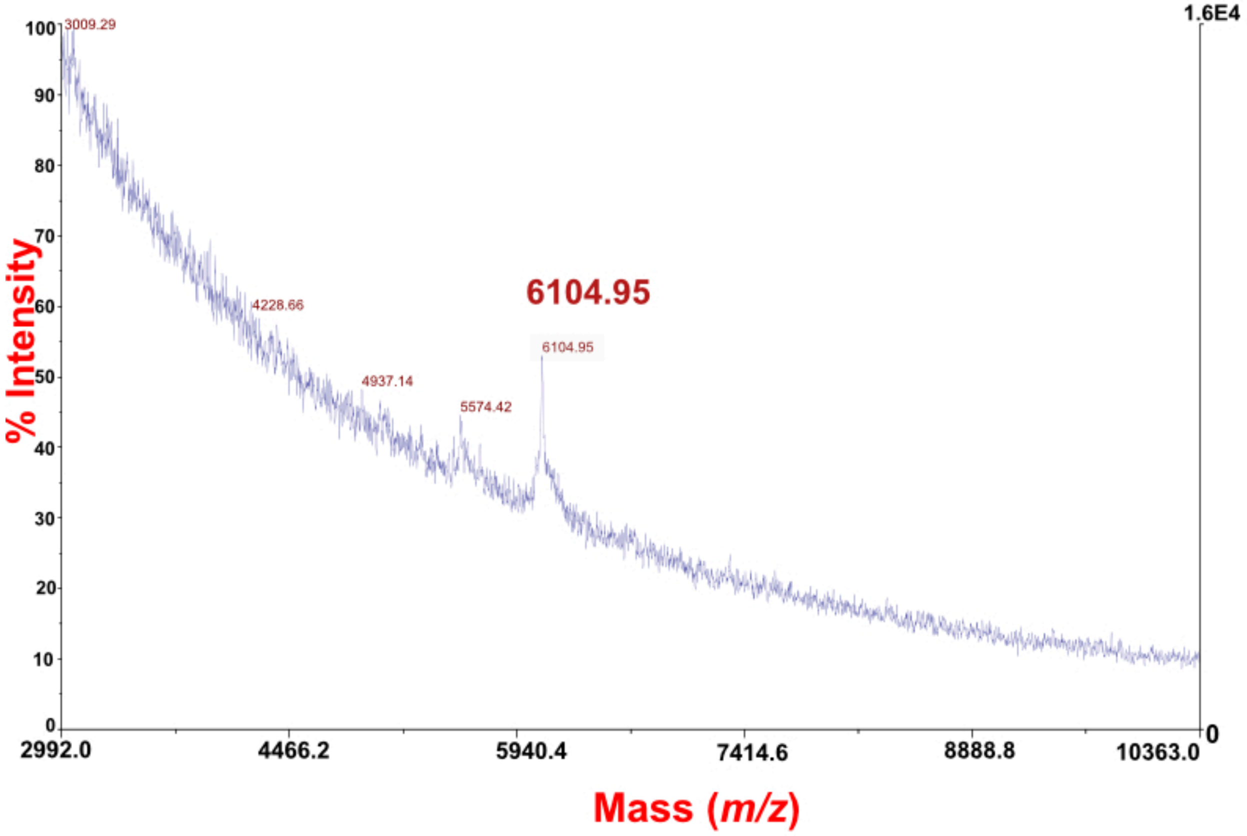


**Figure S10.** MALDI-MS spectrum [%intensity *vs.* Mass (*m/z*)] for pHrodo Red conjugate of Aβ (RODO-Aβ$^{pH}$) corresponding to *m/z* of 6104.95 and 5574.42. These *m/z* peaks are emerging from the conjugation of pHrodo Red (molecular weight of reactive fragment ~535) with an amine functional group of $Aβ_{1-42}$.

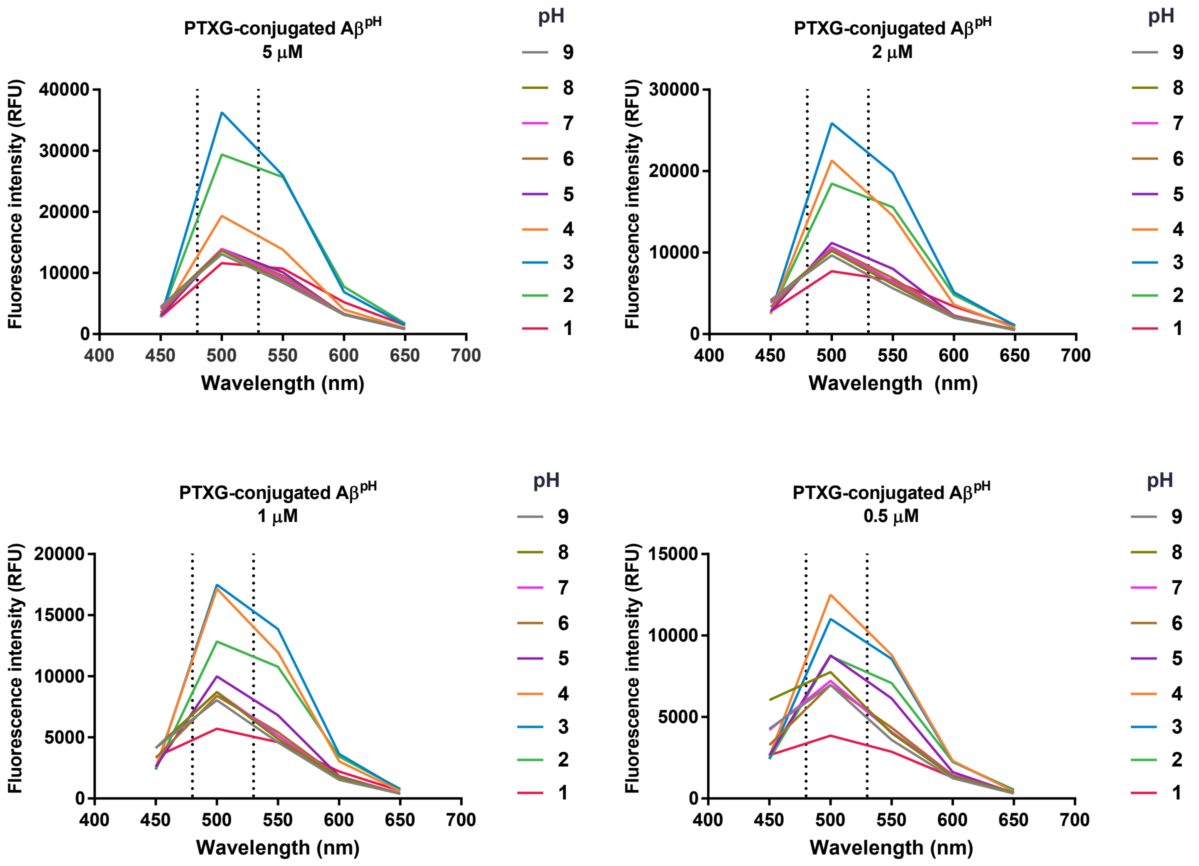


**Figure S11. pH-dependent emission spectra of Protonex Green conjugated Aβ at various concentrations.** The emission spectra of the PTXG-Aβ$^{pH}$ reporter showing maximum fluorescence between acidic pH of 2.0 to 4.0 and limited fluorescence at the physiological pH of 7.4. Dotted lines highlight the region of maximum emission. Legend shows color code for pH 1 to 9.

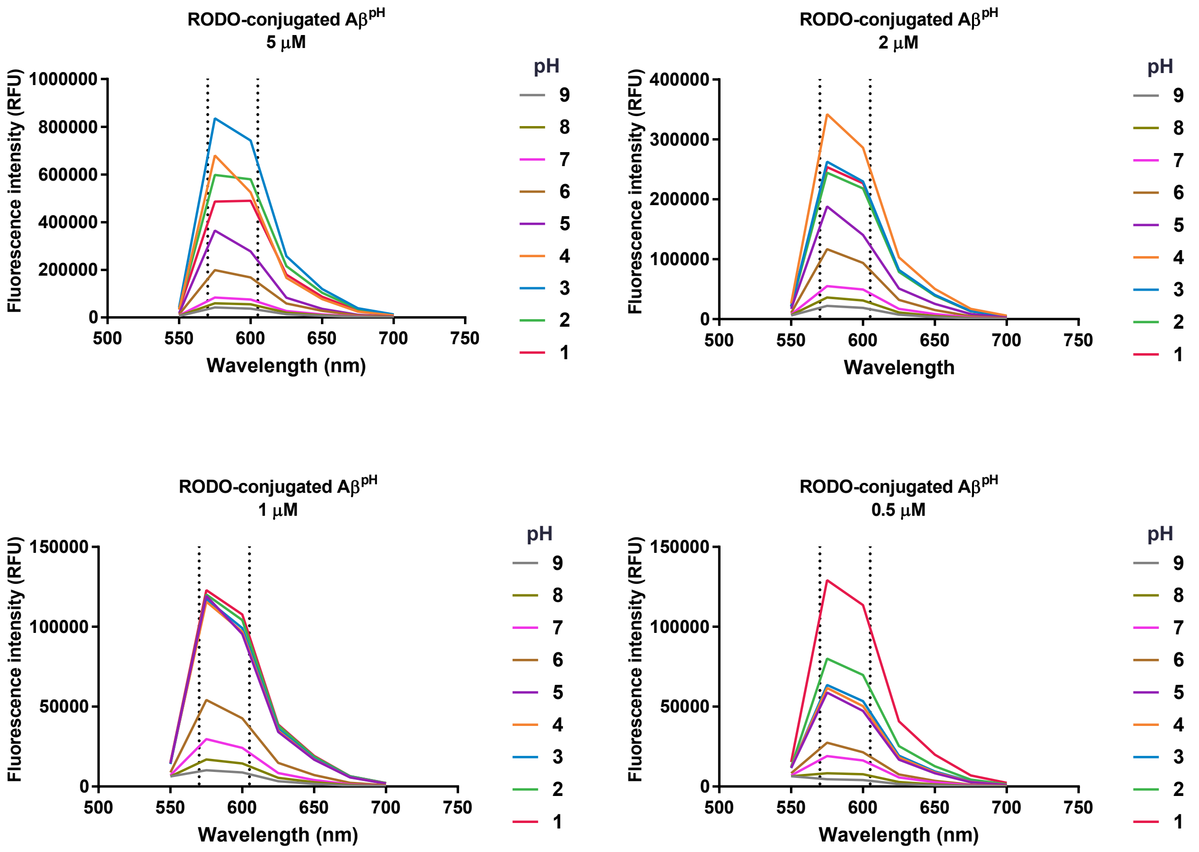


**Figure S12. pH-dependent emission spectra of pHrodo Red conjugated Aβ at various concentrations.** The emission spectra of the RODO-Aβ$^{pH}$ reporter showing maximum fluorescence over a large pH range of 1.0 to 5.0 along with differences between concentrations. Dotted lines highlight the region of maximum emission. Legend shows color code for pH 1 to 9.

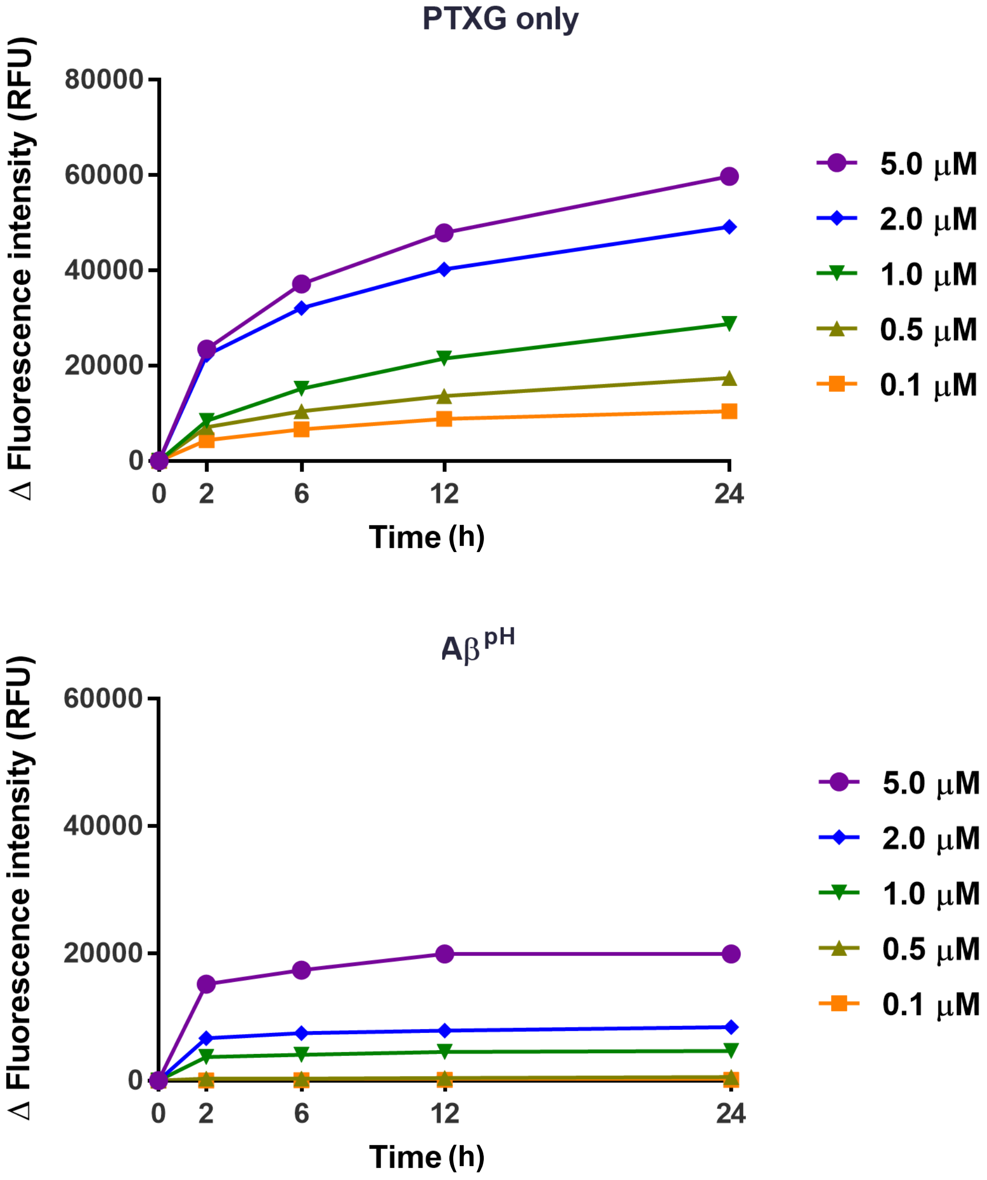


**Figure S13. Concentration-dependent response of PTXG and PTXG-conjugated Aβ (Aβ$^{pH}$) at acidic pH over time.** Fluorescence intensity of the PTXG-conjugated Aβ (Aβ$^{pH}$, bottom plot) in acidic condition of pH 3.0 over a 24 h period. Top plot shows the fluorescence intensity of PTXG alone.

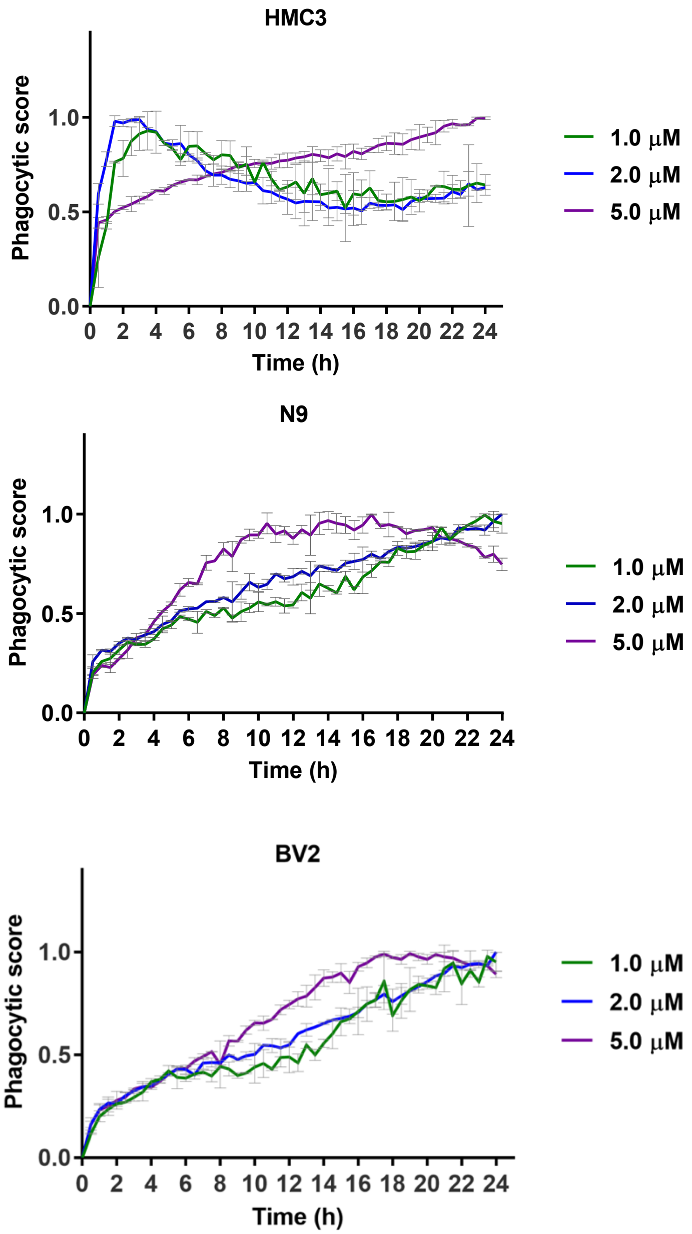


**Figure S14. Change in Phagocytic Score over time.** For human microglia (HMC3, top), 5 $\mu$M of A$\beta^{pH}$ leads to a gradual increase in phagocytic score over time. At lower concentrations of 1 and 2 $\mu$M A$\beta^{pH}$, increased phagocytic score is observed peaking between 2h to 6 h indicating maximum florescence intensity over 24 hours compared to starting (t=0) time point. This is followed by progressive decrease in phagocytic score indicating decreased uptake or increased degradation compared to t=0 time point and ultimately flattening out after 14 hours. In mouse N9 (center) and BV2 (bottom) microglia, higher A$\beta^{pH}$ concentration (5 $\mu$M) results in higher phagocytic score over time peaking at 12-16 hours for N9 and 16-20 hours for BV2 compared to no peaks at 1 and 2 $\mu$M A$\beta^{pH}$ treatment.

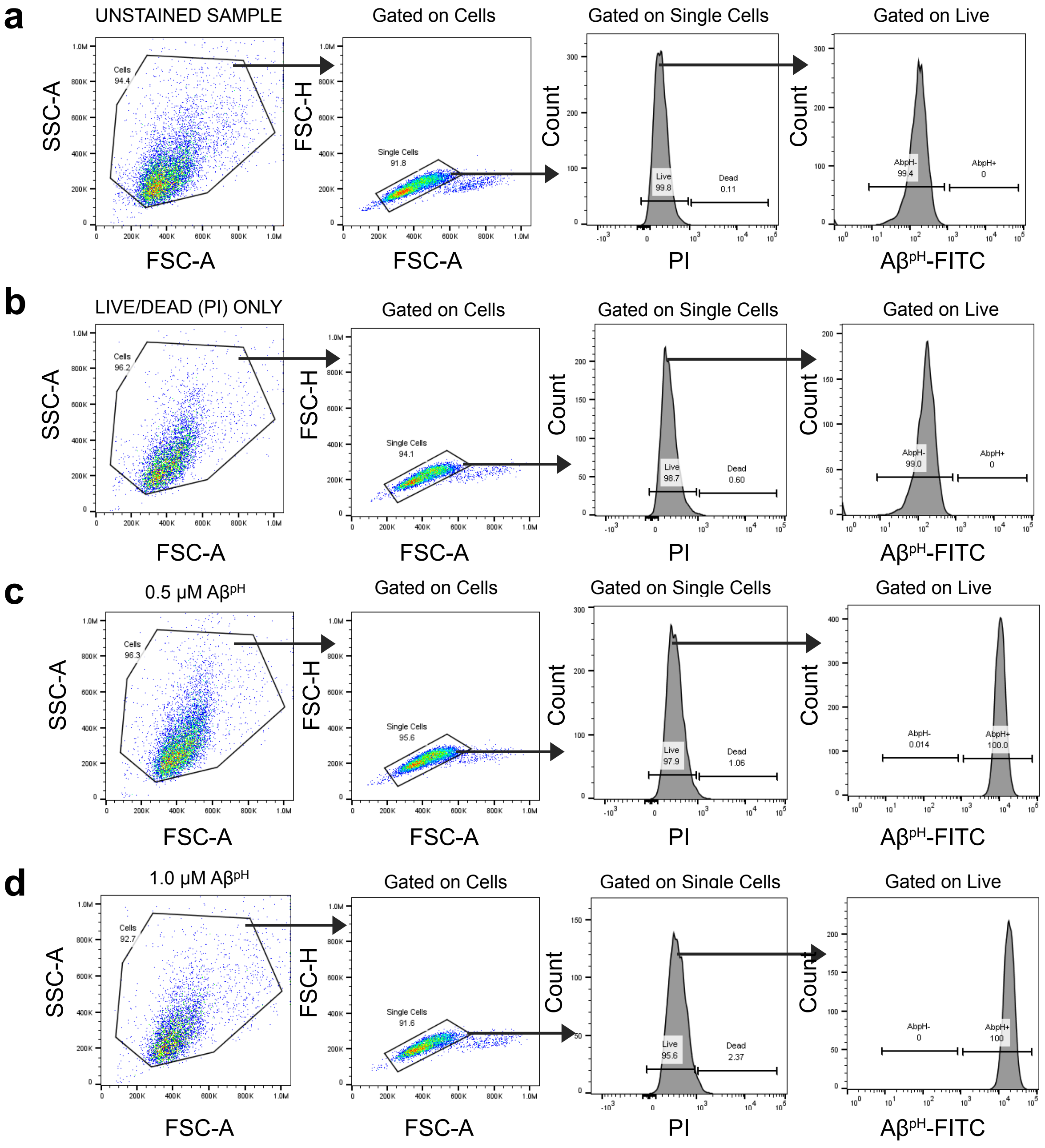


**Figure S15.** Gating strategy for flow cytometry analysis of $A\beta^{pH}$ by BV2 microglia. **a.** Unstained sample i.e. cells only. **b.** Cells with live/dead stain only. Propidium iodide (PI) was used as a live/dead stain (indicating dead cells). **c.** Cells treated with 0.5 μM $A\beta^{pH}$ for 1 h. **d.** Cells treated with 1.0 μM $A\beta^{pH}$ for 1 h.

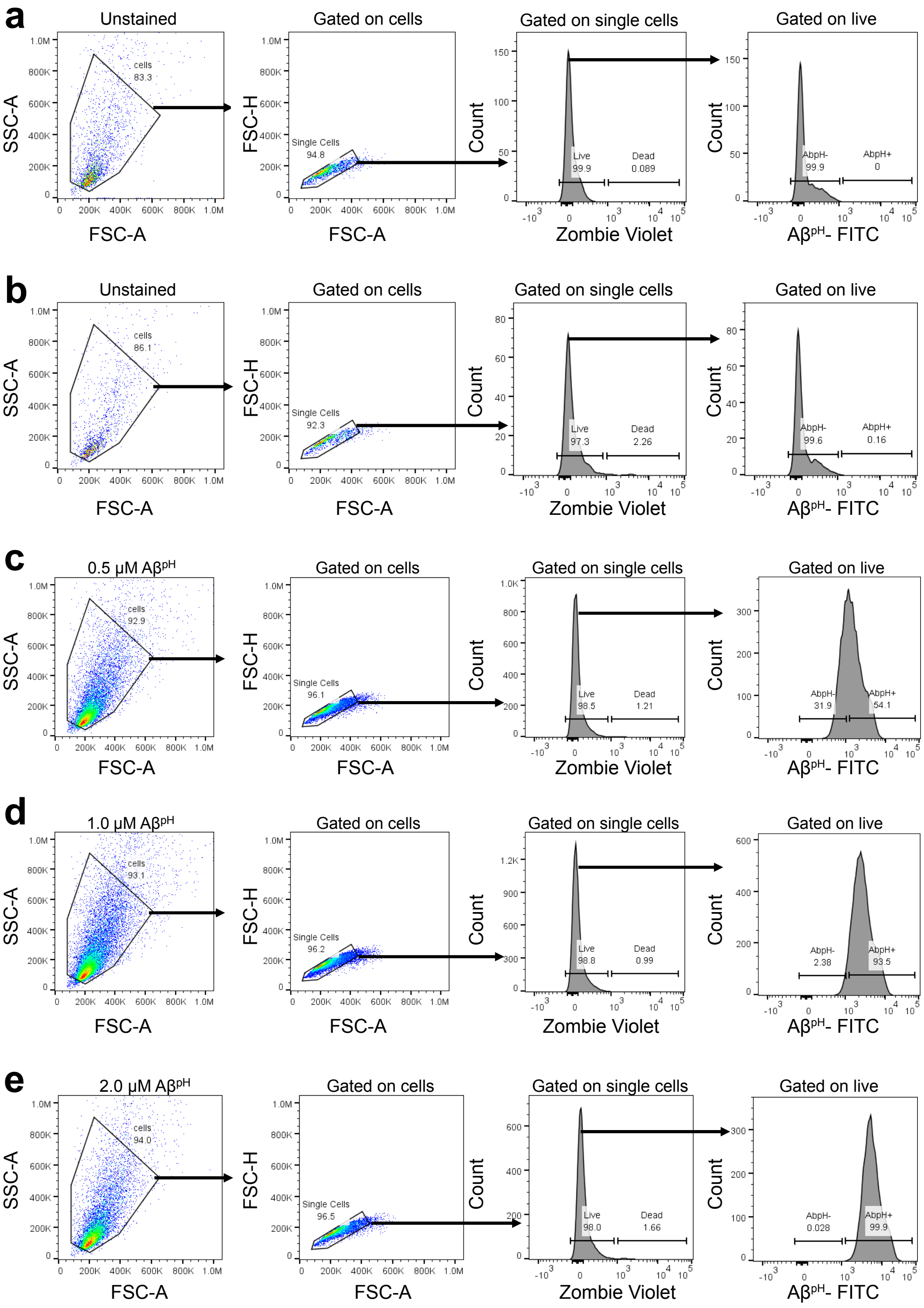


**Figure S16.** Gating strategy for flow cytometry analysis of $A\beta^{pH}$ by primary microglia. **a.** Unstained sample i.e. cells only. **b.** Cells with live/dead stain only. Zombie violet was used as a live/dead stain (indicating dead cells). Cells were treated with **c.** 0.5 μM, **d.** 1.0 μM, and **e.** 2.0 μM $A\beta^{pH}$ for 1 h.

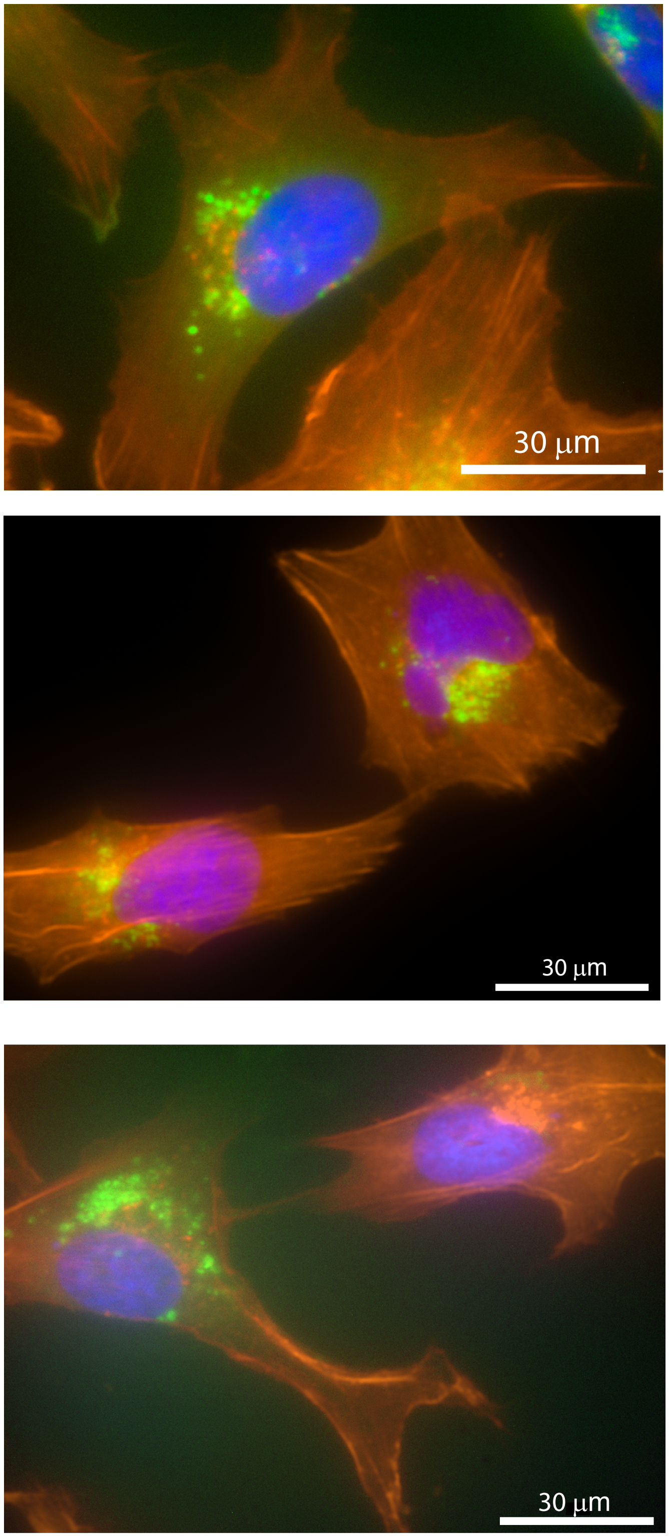


**Figure S17.** Imaging phagocytosis of A$\beta^{pH}$ (green) in fixed HMC3 cells stained with phalloidin (red) and DAPI (blue).

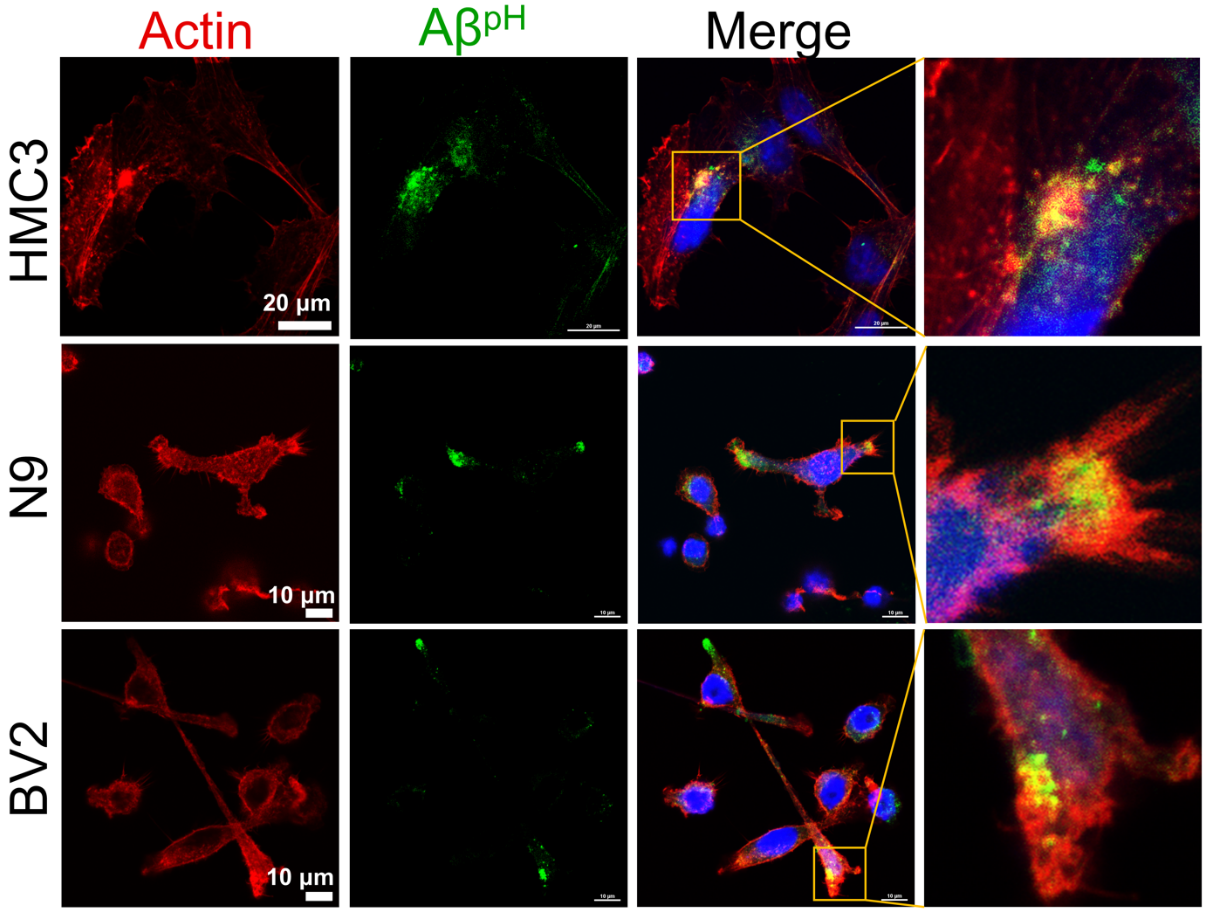


**Figure S18.** Confocal images of fixed HMC3, N9, and BV2 cells showing the uptake of $A\beta^{pH}$ (green). Cells are stained for actin (red) and nuclei (blue) and retain the green $A\beta^{pH}$ signal after fixing.

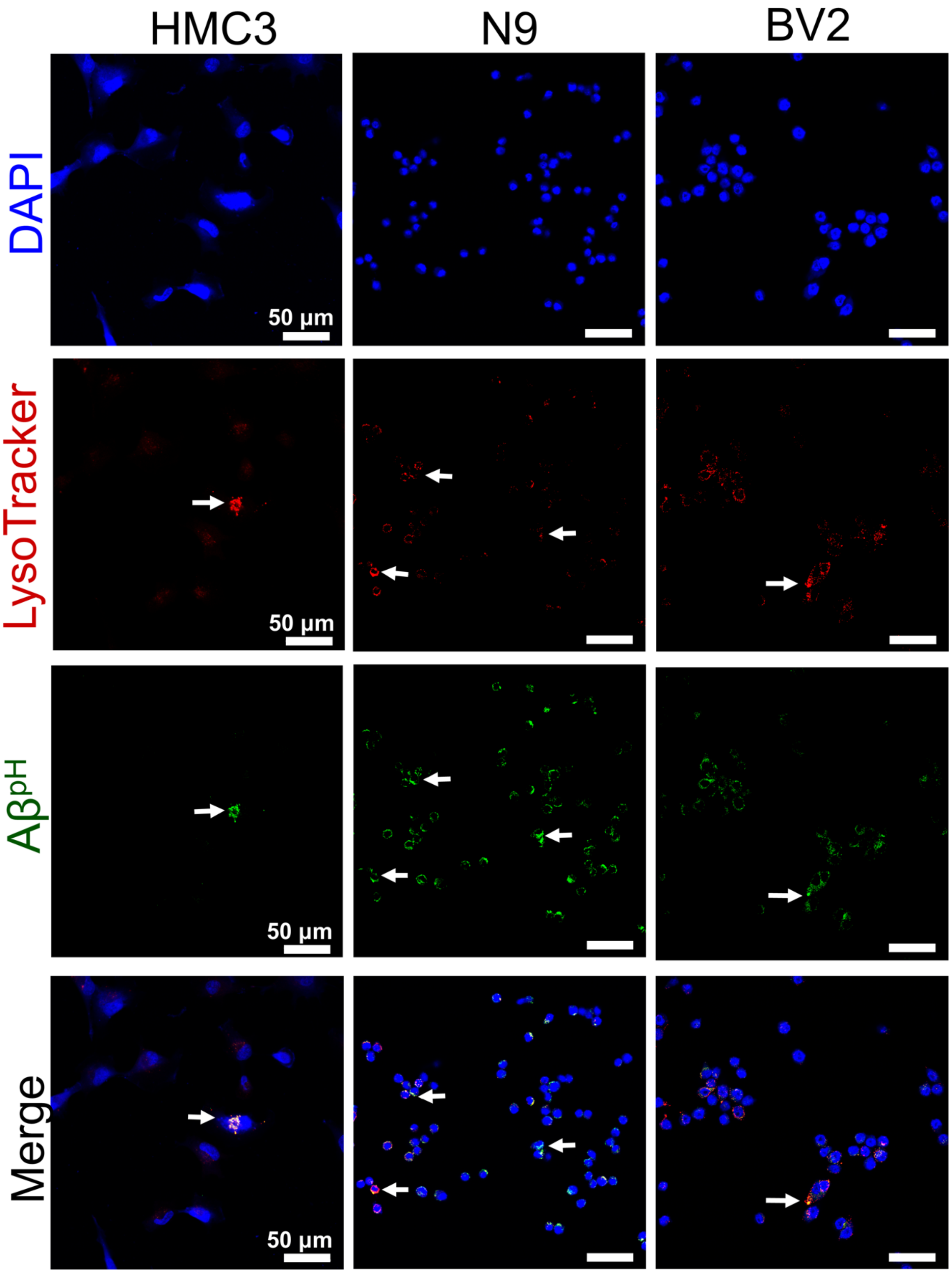


**Figure S19.** Confocal images of fixed HMC3, N9, and BV2 cells showing the uptake of Aβ$^{pH}$ (green) as indicated by the white arrows. Cells are stained for acidic intracellular organelles (LysoTracker, red) and nuclei (DAPI, blue).

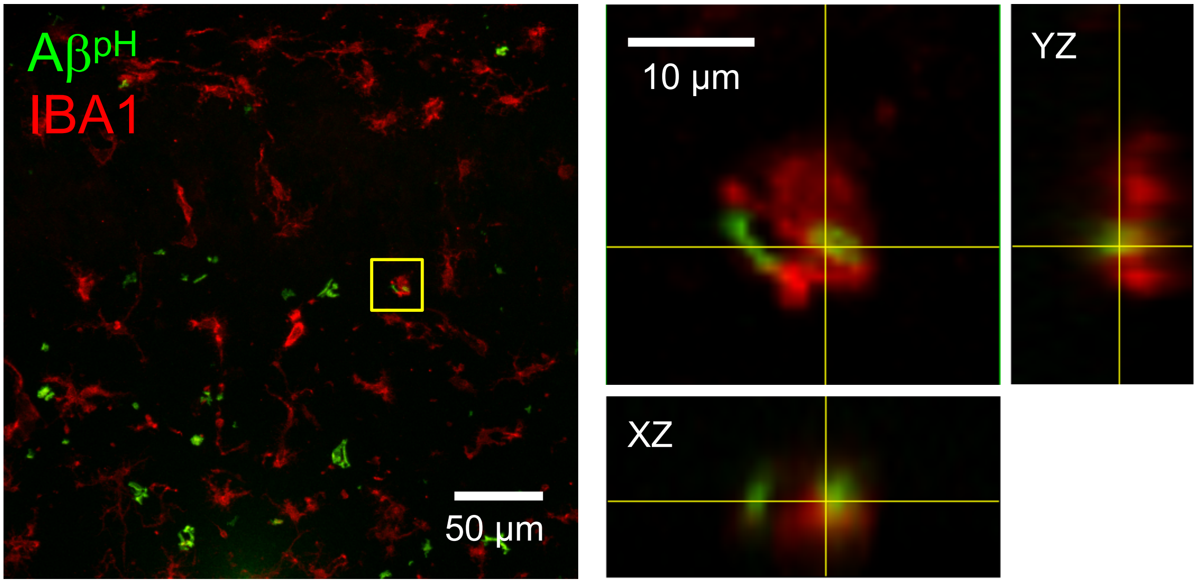


**Figure S20.** Uptake of $A\beta^{pH}$ (green) by $IBA1^{+}$ microglia (red) in acute hippocampal slices from a P12 rat. The $A\beta^{pH}$ is taken up by microglia shown by overlap of green and red florescence (images on right). Green florescence of $A\beta^{pH}$ outside microglial cells (red) indicates uptake into cells other than microglia.
